# Motif distribution in genomes gives insights into gene clustering and co-regulation

**DOI:** 10.1101/2024.09.18.613605

**Authors:** Atreyi Chakraborty, Sumant Chopde, M.S Madhusudhan

## Abstract

We read the genome as proteins in the cell would - by studying the distributions of 5-6 base motifs of DNA in the whole genome or smaller stretches such as parts of, or whole chromosomes. This led us to some interesting findings about motif clustering and chromosome organisation. It is quite clear that the motif distribution in genomes is not random at the length scales we examined: 1kbps to entire chromosomes. The observed to expected (OE) ratios of motif distributions show strong correlations in pairs of chromosomes that are susceptible to translocations. With the aid of examples, we suggest that similarity in motif distributions in promoter regions of genes could imply co-regulation. A simple extension of this idea empowers us with the ability to construct gene regulatory networks. Further, we could make inferences about the spatial proximity of genomic fragments using these motif distributions. Spatially proximal regions, as deduced by Hi-C or pcHi-C, were ∼3.5 times more likely to have their motif distributions correlated than non-proximal regions. These correlations had strong contributions from the CTCF protein recognizing motifs which are known markers of TADs. In general, correlating genomic regions by motif distribution comparisons alone is rife with functional information.

**GRAPHICAL ABSTRACT:** 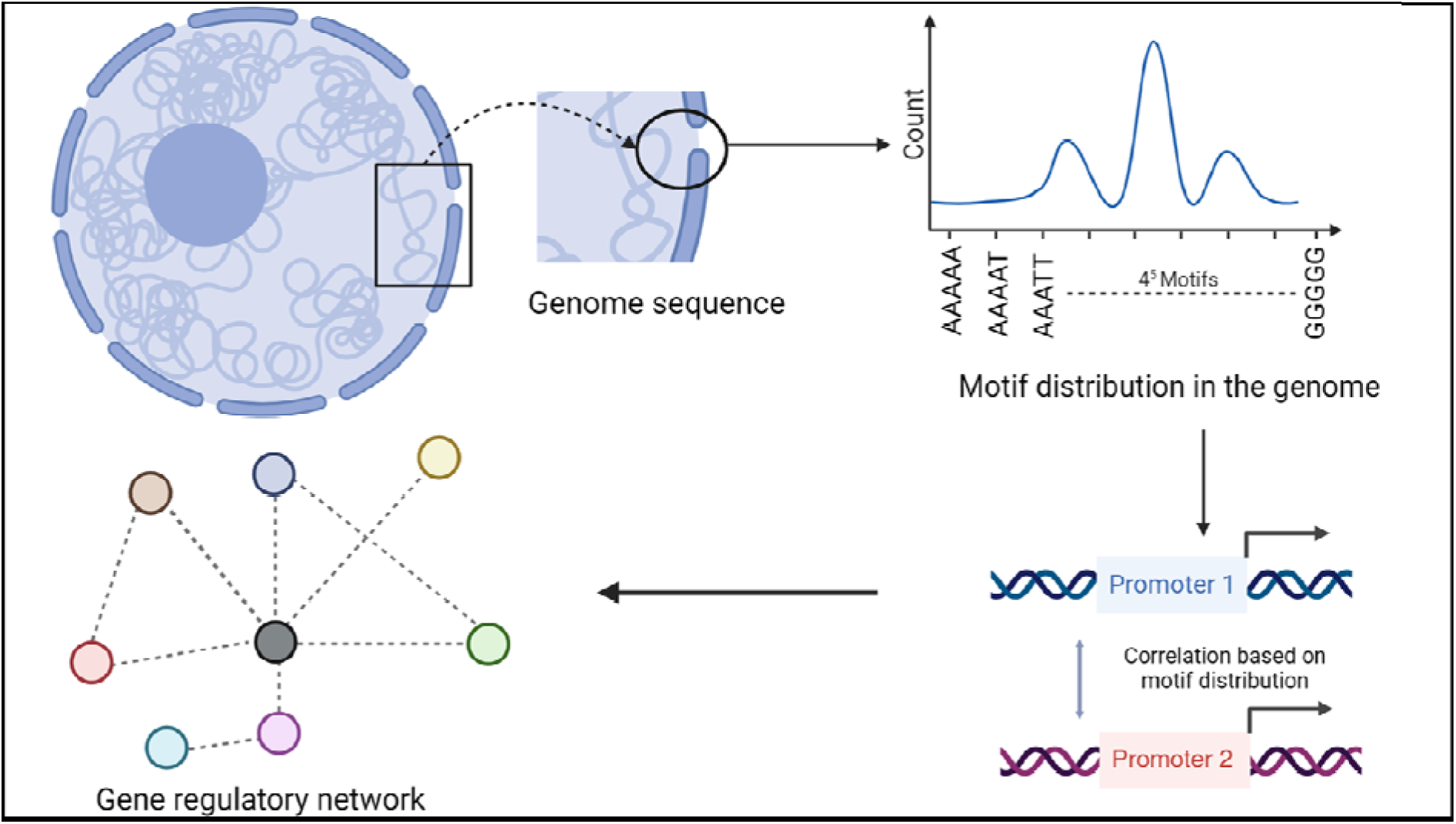

## INTRODUCTION

The human nuclear genome comprises a vast sequence of 3.2 billion base pairs of DNA, organized into 23 or 24 distinct chromosomes. This sequence serves as the fundamental code that underpins the operation of the intricate biological machinery within the cell. Based on the T2T-CHM13 reference ^1^, a comprehensive annotation effort has identified a total of 63,494 genes within the human genome. This count includes both protein-coding genes, pseudogenes, hypothetical genes and functional RNA (lncRNA, miRNA etc.) genes^2^. Genic regions are however only a small fraction of the genome, which predominantly comprises non-coding regions^3–5^. These non-coding parts of the genome harbour an abundance of repetitive DNA, such as transposable elements^6^ and contain valuable information on disease-causing mutations^7^, genetic variations ^8,9^ and evolutionary conservation ^10,2^. Analyzing genomes for all of the characteristics mentioned above usually involves aligning sequence stretches to one another. In this study, we offer an alternative yet simple method to ‘read’ the genome.

The regulation of the transcriptome inside the nucleus is complex^11^. For instance, genes and their corresponding regulators may be sequentially distant but spatially proximal ^12,13, 14^ as the regulation is coordinated by proteins. Different techniques are used to investigate this spatial organization, protein-genome interaction and chromatin accessibility such as 3C techniques ^15^, Hi-C ^9,16^, Dam ID ^17,18^, Chip-Seq^19,20^, ATAC Seq^21,22^, super-resolution imaging^23^ and integrative genome modelling^24,25^.

The resolution of the information from the techniques mentioned above is usually in the range of 10-100 kilo-base pairs(kbps). Investigating genome stretches of size 10-100 kbps, especially with the intent of gaining insights into genome organization/function poses several difficulties. Firstly, the 10-100 kbps size range is very low resolution, given that DNA binding proteins engage directly with just a few base pairs. Secondly, even if one were to look for sequence conservation and/or patterns, 10-100kbps is a significantly large stretch compounded by the fact that the DNA alphabet consists of only 4 letters. Sequence alignments of such large stretches are often poor indicators of functional conservation. Even two randomly chosen sequence stretches of ∼10-100kbps could be ∼25% identical. Therefore, identifying and comparing sequence patterns necessitates a focus on shorter sequence sizes. Alignment-free methods, such as k-mer or word frequency estimation, offer alternatives where sequences are processed in moving windows of a specified word length^26,27^. Notably, this approach does not hinge on positional information of bases in sequence. However, it proves valuable for comparing lengthy sequences, allowing the calculation of word density to determine the similarity or dissimilarity between two sequences^28,29^. This method provides a practical means for assessing sequence patterns without relying on extensive alignments, particularly suited for analysing large genomic regions. In this study, we used a modification of the k-mer method to compare genome segments.

Binding patches on proteins typically recognize 5-6 bases of DNA^30^. Our observation from all known DNA-bound protein complexes in the PDB is that at the points where proteins contact DNA specifically, the interactions only span 5-6 nucleotides. However, there are several reports of larger DNA sequence motifs recognized by proteins^31,32^. These happen in cases where the protein could oligomerize and hence recognize several, seemingly contiguous patches of DNA that are larger than 6 nucleotides. It could also happen when DNA wraps around a protein making contacts at different faces (supplementary figure 1). Our observation is that for every contact only 5-6 nucleotides are involved in base specific recognition such as hydrogen bonding. Hence, we focussed on sampling motifs of 5-6 bases of DNA and read the genome at that resolution. We used the term motif to represent an n-mer assuming that all n-mers are potential motifs for some cognate protein(s). In this study, we first analysed motif distributions of different sizes (from 2mers to 6-mers). We then compared this to a randomised genome to show that patterns in real genomes are distinct. We examined patterns of motif distributions in whole genomes/chromosomes and in smaller regions such as centromeres and gene promoters. We correlated pattern distributions in gene promoter sites and established relationships between genes that are likely to be co-regulated. Sometimes when protein interact with DNA, the DNA could bend or alter its conformation from the regular B-form^33,34^. That notwithstanding, in this study we are assuming that a particular DNA motif when binding to its cognate protein would always undergo the same recognition shape change.

We corroborated our findings, using three case studies. In the first case study, we selected three genes, LPHN1(ADGRL1), CDK9 and TRIM8, that exhibited high correlation based on our motif abundance analysis and compared them to a network map using NetworkAnalyst ^35,36^. We identified two common transcription factors to all three genes, thus validating not just our method of constructing gene networks but also discovering the functional importance of such connections. For the second case study, we focused on a larger list of 19 genes coregulated by the transcription factor(s) Jun/Fos. The analysis here showed that we could predict differential gene regulation. In the final case study, we investigated whether gene co-regulation implied colocalization by comparing our results to Hi-C^37^ and promoter capture Hi-C^38^ data. We found a strong correlation between our scoring and the spatial positioning of genome segments, reinforcing the significance of our methods.

## MATERIAL AND METHODS

Data of genomic sequence(s) and the gene annotation, for all calculations was obtained from NCBI RefSeq Human Genome assembly T2T-CHM13v2.0. ^39,40^

### 1. Motif generation and distribution metric scoring

To assess the enrichment of specific motifs within certain regions, we employed the observed-to-expected ratio (OE ratio). The OE ratio is a measure of whether a feature is over or underrepresented in a given dataset. It is calculated by normalising the observed frequency of the feature by the expected frequency based on the probability of occurrence. For a sequence stretch of base pairs, we can ‘read’ it using a window (motif size) of *n* base pairs. The number of times a motif occurs in the sequence stretch is recorded as the observed count (O). The expected count (E) of any motif is (N - k + 1) *∏* f^nX^, where *nx* is the number of times a particular base occurs in a motif and whose probability is *f*_x_ and *N* is the size of the chromosome. The OE score is thus a normalized metric taking into account the AT/GC richness of a chromosome. Given that there are 4 bases (A/T/G/C), the number of motifs of size *k*, which we refer to as k-mer count, would be 4^k^. Here we average the OE values for motifs and their reverse complements. These averaged OE ratios taken together are called the motif vector. We recorded the observed count of motifs and then computed the OE ratio for the following sequence stretches:

- **OE _whole_** : Here the sequence stretch is the whole chromosome.
- **OE _nkbps_** : Chromosomes are divided into bins of n kbps. The last bin of any chromosome may contain less than n kbps, while all other bins are exactly n kbps in length.
- **OE _centromere_** : Only the centromeric regions were considered. The boundaries of centromeric regions in the different chromosomes were taken as defined in the NCBI genome data viewer and UCSC genome browser ^40,41^ (supplementary table 1).
- **OE _promoter_** : We sampled 3 different sequence stretches in the 5’-UTR that we define as promoter proximal control regions. For simplicity we refer to this as promoter regions in the rest of the text. These were 1 kbps, 2 kbps and 6 kbps upstream of the gene transcription start site and denoted as OE_1kbps_kmer_, OE_2kbps_kmer_ and OE_6kbps_kmer_, where motif size k=5 or 6. We obtained the gene start and stop coordinates from the NCBI RefSeq gene annotation ^40,41^.
- **OE _random_** : For different computations (as expounded in the results), it was necessary to scramble a sequence stretch and rerecord the observed count. To do this, the sequences of entire chromosomes were shuffled (randomised). The procedure for randomization is mentioned in methods section 3.

Note that in all these computations, the observed count is taken from the different sequence stretches while the expected count is computed based on chromosome size (N) and is the same across all computations (for a particular chromosome).

### 2. Correlating motif vectors

The correlation between two motif vectors, X and Y, is computed as:

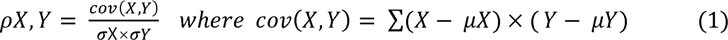

Where cov(X, Y) is the covariance, (µX and µY) and (oX and oY) are the means and standard deviations of OEs of motif vectors X and Y respectively.

### 3. Generating random genome

To randomly generate DNA sequences of different chromosomes, we used the random module of python3 and ensured weighted random nucleotide selections. This procedure generates random sequences of chromosomes of the same size and base probability as the wild type. The random genome constructed thus has the same nucleotide composition as the original, just scrambled in sequence.

### 4. Finding Hi-C contacts with genes

Hi-C data were obtained from GEO accession number GSE18215 (in .mcool format), where chromosomes were divided into uniform bins of different sizes. We chose to use data from the bins of the smallest size (highest resolution), 10 kbps. We extracted pairs of 10kbps bins from the data where cross links were recorded between chromosomes 18 and 19.

For the promoter capture Hi-C (pcHi-C), we extracted the chromosome contact start-stop coordinates from the .bedpe files from GEO accession number: GSM1704495. From the NCBI gene annotation for the corresponding reference assembly used in mapping the contacts, we obtained the gene coordinates that overlap with the contact regions between the two chromosomes. For the gene pairs that fall in the region of contacts, we obtained the promoter correlations according to our motif vector calculations at the different aforementioned promoter sizes.

## RESULTS

### 1. Motif Preferences as assessed by the OE ratio

We had previously established that protein binding stretches in genomic DNA span 5-6 nucleotides ^30^ (supplementary figure 1). So, in this study, we have read the genome at this length scale, i.e., at biologically relevant DNA motif sizes. To begin with, we read the genome with motif sizes of 2 - 6 nucleotides. The aim here was to check the effective size of motifs that would confer specificity to DNA-protein binding. We noticed that as the motif size increased from 2 to 6, the number of motifs with high OE_whole_ ratio values increased (figure 1a and supplementary figure 2). While the trend is monotonic, we did not explore beyond 6-mers as this is the optimal protein recognition size and we believe that all specificity discriminations are likely to occur at this length. The inference here is that the larger the variation in the OE_whole_ ratio values, the more pronounced the patterns of occurrences of these motifs. This in turn has implications on how different proteins would engage with different parts of the genome.

**Figure 1:**
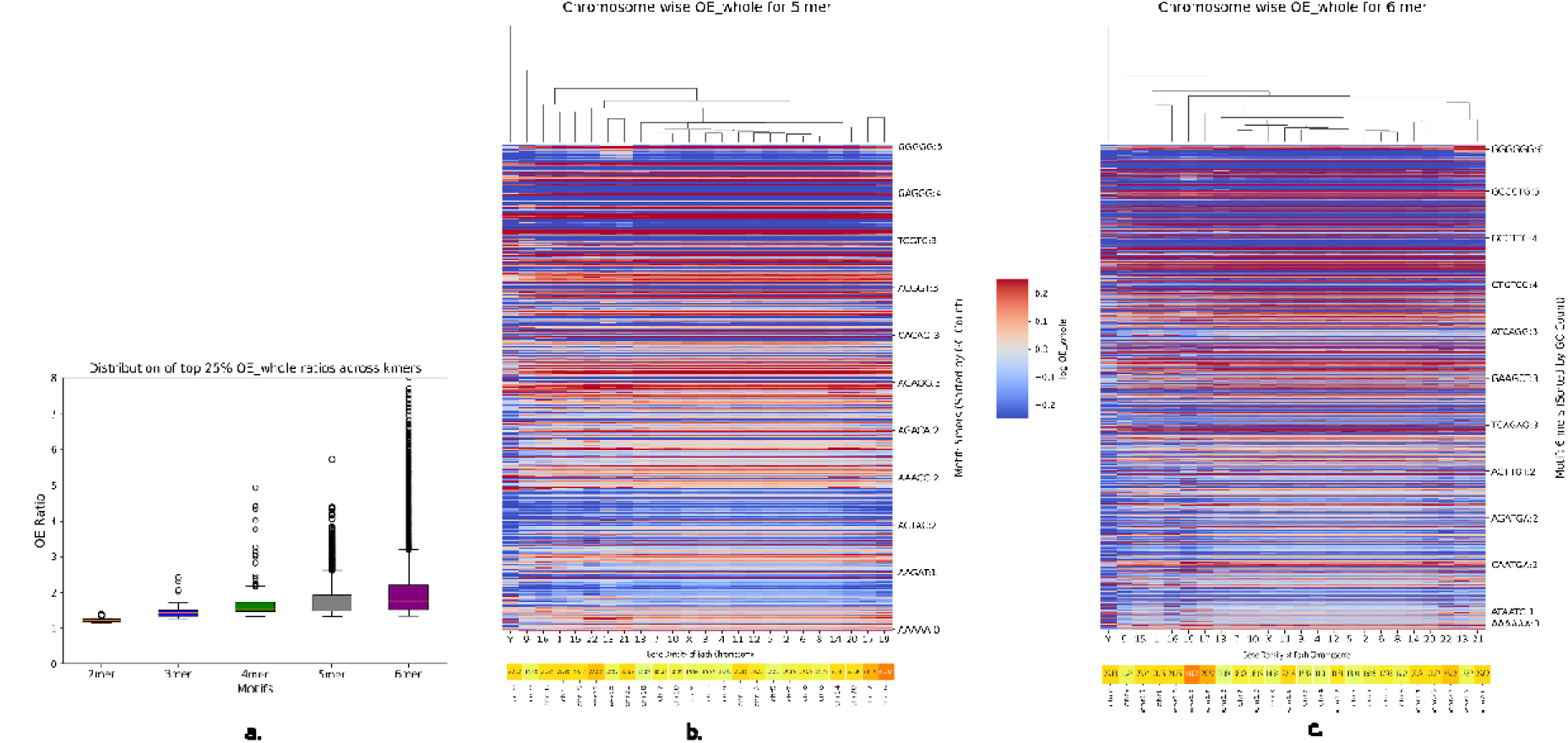
The distribution of OE_whole_ ratios for motif size 2-6 (a). The OE ratios are limited to the range from 0 to 8 and motifs of sizes 2,3,4,5 and 6 are represented in different colours. Heat maps of OE_whole_ ratios for motif size 5-6 (panels b-c respectively) across the different chromosomes shown in log scale. The log(OE) ratios are coloured red through to blue (colour legend) where red and blue indicate values of > 1 and < 1 respectively. The darker the shade of the red (or blue) the higher (or lower) the ratio. The 512 and 2080 motifs for 5-mers and 6 mers respectively are arranged according to GC content, which increases going from bottom to top. In panels b and c, only a few representative motifs (and their reverse complements) are labelled. Shown alongside the motif labels are their GC contents. The dendrogram represents the clustering of chromosomes based on the similarity in motif distribution. A colour bar of gene density is shown at the bottom of panels b and c. The denser is the gene content the redder is the colour bar.

For the rest of this study, we concentrate on 5- and 6-mer motifs. We next looked at the OE_whole_ ratio of 5- and 6-mers for all individual chromosomes (figures 1b and c). While considering OE ratios we averaged the values of a motif and that of its reverse complement as we are looking at double stranded DNA. Hence, we got 512 different motifs of 5-mers and 2080 motifs of 6-mers. (The number of 6-mer motifs and their reverse complement pairs is not exactly half of all possible motifs (4,096) because palindromic motifs are identical to their reverse complements). There are two appreciable patterns to discern here – a) Certain motifs are less abundant in the whole genome while others are present in large abundance, as inferred from their OE_whole_ ratio values. For many motifs, these patterns of abundance hold across the different chromosomes. For instance, the 6-mer motif CCCAGC has high abundance in all chromosomes. The CCCAGCAG is a binding site for the zinc finger, ZNF143^42^, a protein involved in 3D genome construction. b) There are variations in the OE_whole_ ratio values for the same motif across chromosomes. We clustered the chromosomes based on their motif distribution similarity (represented as dendrograms in figures 1b and 1c). The gene density (number of genes per Mbps of a chromosome) of individual chromosomes is independent of the chromosome clustering based on the motif OE ratios (figure 1b and 1c and supplementary figure 3). We find that the clustering of chromosomes when considering 5-mers or 6-mers is similar (with minor rearrangements). Chromosomes 2- 8,10-12,14,18, 20 and X are all more closely related to one another than to the others. Among the others, (17,19), (1,15,16), and (13,21,22) form smaller sub-clusters of similar chromosomes. Chromosomes 9 and Y are the two most distinct ones and bear the least resemblance to the others (see supplementary figure 4a for pairwise correlations).

On closer inspection, it is clear that each chromosome has its unique pattern of motif abundance. For instance, it is plain to see that the 5-mer and 6-mer motif abundances of chromosome 9 and Y are starkly different from the other chromosomes (figures 1b and c and supplementary figure 4a). Chromosome 9 is known for its significant structural variability, housing the largest autosomal block of heterochromatin^43^. Within this region, gene locations exhibit a distinctive pattern, with individual genes scattered among stretches of repeated sequences^44,45^. The case is similar to chromosome Y, which is known to have the maximum occurrences or repeat regions^46^. These results suggest the possibility that different parts of the genome are recognized differentially, presumably, by different proteins^47^. These recognition events in turn could differentially regulate gene expression and other molecular functions^48^.

### 2. The distribution of motifs in chromosomes is non-random

From the results above it is clear that motifs are present with different abundances in the different chromosomes. What is also clear is that while some broad conservation patterns may exist, the motif distribution in individual chromosomes is distinct. Within each chromosome, sub-regions have different distributions of the motifs, as seen in the 6-mer motif distribution along 100kbps stretches of chromosome 18 (figure 2a). Details of the distribution of 5- and 6-mer motifs for all chromosomes are presented in supplementary data. Chromosome 18 has ∼80.3Mbps, which were divided into regions of 100kbps. We retained the same composition of bases and randomly scrambled the sequence of the whole chromosome (figure 2b). The distinct patterns visible in the real distribution (OE_100kbps_) are no longer apparent in the scrambled sequence (OE_random_). We did the randomization multiple times) and in each such attempt, the distinctness of the pattern of the real chromosome (in 100kbps bins) was absent (supplementary figure 5 and 6). The only discernible pattern in the randomised chromosome is the similarities of motif distributions of distinct GC content. It is interesting to note that the bin wise motif distribution is distinctly different in regions that correspond to the centromere (supplementary figure 5). The centromeres can be easily distinguished in each of the chromosomes. This can be attributed to the satellite repeat regions in the centromeres^49^.

**Figure 2:**
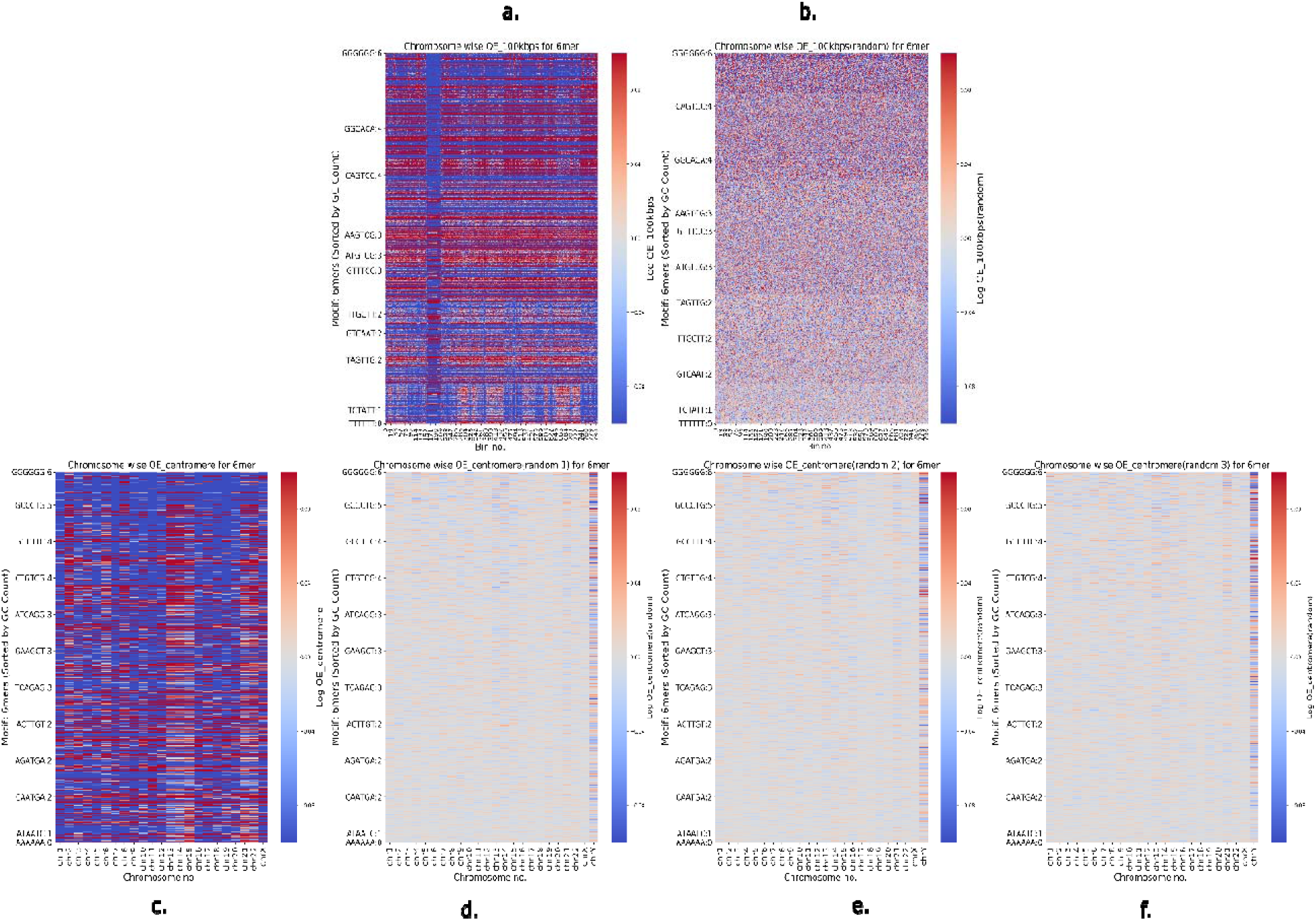
Panels a and b show the OE_100kb_ and OE_random_ heat maps for chromosome 18 in log scale respectively. Heat maps of the log values of motif OE_centomere_ ratios of 6-mers for centromeric regions in all chromosomes (panel c). The randomised sequences of the same centromeres are shown in d-f. As in figure 1, only representative motifs and their reverse complements are labelled and their GC content is shown alongside The colour coding and the arrangement of motifs is the same as in figure 1.

As mentioned above, the pattern of motif abundances varied not just across chromosomes but also within chromosomes. While this is evident for 100 kbps bins across the length of chromosome 18 (OE_100kbps_ distribution in figure 2a), we next investigated centromere regions on all chromosomes. The boundaries of centromeric regions of the different chromosomes were taken as defined in the NCBI genome data viewer and UCSC genome browser^40,41^(supplementary table1). Here again, it is clear that the centromeric regions display distinct motif distributions. The OE_centromere_ distributions across the different chromosomes of the genome are different (figure 2c). For instance, the acrocentric chromosomes 13,14,15,21 and 22 have a distinctly different OE_centromere_ pattern (figure 2c and supplementary figure 4b). It is known that the short arms of these acrocentric chromosomes are abundant with segmental duplications^50^, which could be the explanation for our observation.

We next compared the centromere regions to the same regions in the randomly scrambled chromosome sequences (figures 2d-f). Here too, the difference between the motif distribution patterns in the real centromere (OE_centromere_) and the centromere from the scrambled chromosome is clear – the centromere from the scrambled sequences are not distinct across chromosomes. We triplicated this result for confidence (figures 2d-f).

We have established that motif distributions in chromosomes are not random and that such distributions are also different in different sequence segments within each chromosome. This gives further credence to the point made earlier that the distinct motif distributions ensure differential binding by the different protein partners.

### 3. Gene correlations deduced from promoter motif distributions

In the next few sections, we concentrate on the 63,494 genic regions, as identified by the T2T CHM13 reference assembly. We variously defined promoter proximal control regions (promoters for short) associated with a gene as the regions 1kbps, 2kbps or 6kbps upstream of the gene transcription start site. These promoter sizes were chosen considering a conservative estimate (1kbps), a maximum known effective size (6kbps)^51^ and a value in between (2kbps).

We considered the relative abundances of all motifs (separately for 5-mers and 6-mers) as motif vectors. The coefficients of the vectors are the 512 and 2080 OE_Xkbps_k-mer_ values for k-mers (k= 5 and 6) respectively, where X is the size of the upstream regions considered as promoters. All vs all gene promoter motif vectors were correlated (equation (1)) and the results were collated chromosome-wise (figure 3). Of the ∼4 billion possible gene-gene motif vector correlations, about ∼1 million have correlation coefficients >= 0.9. (the numbers are of a similar order for 6-mers and for different promoter sizes – 2kb and 6kb, see supplementary data of promoter correlations above 0.9). We decided to impose this stringent cut-off to ensure that the genes with correlated promoters are chosen stringently and practically this would mean that we only deal with a small fraction (∼0.25%) of a very large dataset. As the promoter size increases from 1kb to 6kb the baseline of correlations changes while the trends remain the same (supplementary figure 7a-e and supplementary data of promoter correlations above 0.9). The baseline notwithstanding, some features of chromosome-chromosome correlations stand out. Most of the high correlations between gene promoter motif vectors are from within the same chromosome. This is understandable as genes that are functionally related are often found in the same chromosome and in close proximity to one another. On average the number of intra-chromosome gene promoter correlations above 0.9 is ∼920 per chromosome pair (when considering 5-mers with 1kbps promoters). This signifies that on average 920 gene pairs from two different chromosomes are correlated with coefficients greater than 0.9. The maximum number for inter-chromosomal gene promoter correlations is 36,408 between chromosomes 13 and 21, while the minimum is 0 between chromosomes Y with 1 and 19 (supplementary data of promoter correlations above 0.9).

**Figure 3:**
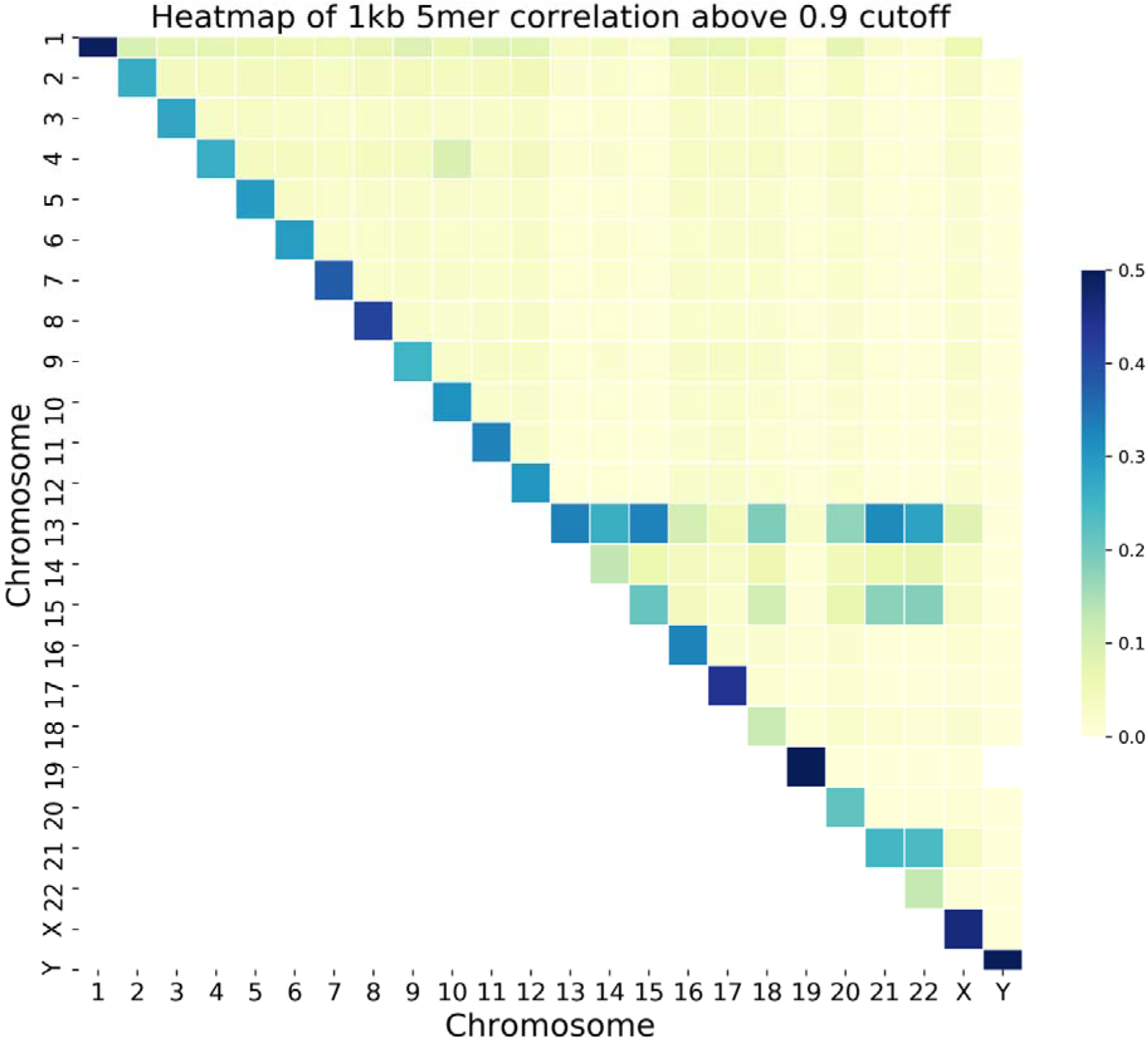
An inter-chromosome motif vector correlation map of frequency of gene pairs with correlation coefficients > 0.9 for 5 mers with promoter sizes of 1kbps. The proportion of correlated genes with a correlation coefficient >= 0.9 are colour coded on a white-yellow-green-blue gradient. The white end represents no correlation while the darker the shade of blue the larger the number of correlated gene pairs.

These correlation data are vast and we are only going to present a figment of it in this study. These data can be interrogated to answer various questions about gene associations. What is of interest to us here are the inter chromosome interactions. There are genes from different chromosomes whose promoters have similar motif vectors. This implies that these genes could be co-regulated. The high correlations, especially ones with coefficients >= 0.9, could be symptomatic of interactions.

Once again the acrocentric chromosomes (13,14,15, 21 and 22) show strong correlations to one another. We found several hundreds of genes in each of the acrocentric chromosomes whose promoter regions were correlated to one another (see supplementary data of promoter correlations above 0.9). Of these, there are 1768, 2216, 2072, 992 and 1310 genes in chromosomes 13, 14, 15, 21 and 22 respectively, that are correlated to at least two promoters from genes of other acrocentric chromosomes. This is interesting as not only do the centromeric regions of these genes show high correlations (section 2 above), but the promoters of genes in these chromosomes show high correlations too. This we believe may be the basis of Robertsonian translocations^52,53^, where the long arms of these chromosomes are joined to one another^52,53^. If the motif abundance is similar in two regions, it is conceivable that recombination could sometimes result in interchanges, resulting in translocations. Ribosomal RNA genes constitute ∼4-13% (supplementary table 2a) of the total genes in chromosomes 13, 14, 15, 21 and 22. Additionally, we also checked what fraction of the total high correlations (Pearson’s correlation coefficient >= 0.9) were contributed by the rRNA gene pairs from both chromosomes. We observed ∼81-92% of the high correlations are contributed by the rRNA genes from both the chromosomes for chromosomes 13,14,15,21 and 22(supplementary table 2b and supplementary data of rRNA gene correlations). Thus, such tandem repeats of rRNA genes constitute the conserved patterns across these chromosomes. Additionally, we compared the similarity in the promoter sequence of the highly correlated genes (Pearson’s correlation coefficient >=0.9) in chromosomes 13 and 14 (chromosomes reported to be involved in translocations) vs chromosomes 18 and 19. 20% of the highly correlated gene pairs between chromosome 13 and chromosome 14 have a sequence identity of at least 80%. However, there are no gene promoter pairs between chromosomes 18-19 that have sequence identity higher than 37% (supplementary figure 8 and supplementary data of identity vs correlations). This suggests that translocations occur in regions where there is strong motif vector correlation as well as high sequence identity.

Another notable gene promoter correlation was between chromosomes 4 and 10 where 1017 (supplementary data of rRNA gene correlations) gene promoter pairs had a correlation coefficient above 0.9 (figure 3). A translocation between these two chromosomes is also known in certain types of leukaemia^54,55^ and Wolf-Hirschhorn syndrome^56^. Unlike acrocentric chromosomes, the gene promoter pairs does not include rRNA genes. However the reason for the translocation could be the local similarity in the motif distribution.

As a control, we repeated the analysis in three pairs of chromosomes for regions 1kb downstream of the gene and also for 1kb regions chosen randomly from the chromosome. In each of these controls, the number of selected regions was the same as the number of gene promoter regions. We selected the following chromosome pairs: (6,7), (12,13) and (11,18). The pair selections were done to include chromosomes of different sizes and gene densities (supplementary figure 3). We obtained the correlations of 5- and 6-mer motif vectors in gene promoters, downstream regions and randomly selected regions.

For the three pairs of chromosomes chosen, there are between 892,167 and 4,552,601 possible gene-gene correlations. Of these, between 1-2% of the promoter correlations are >= 0.5 for these three pairs. Interestingly, the correlation between downstream regions is ∼0.5% in all three pairs (table 1a and supplementary data of promoter vs downstream vs random correlations). This can be partially explained by the fact that in each of the selected chromosomes, ∼20% of the genes overlap with one another (table 1b). It is likely therefore that promoters of one gene could be the downstream region of another and vice versa. Also many closely located genes may be controlled by a single promoter region. All this notwithstanding, there are however a significant number of high correlations between downstream regions that warrant further investigations. The random regions of 1kb show the poorest correlations among all three pairs, accounting for less than ∼0.2% of all correlations. In fact, for the chromosome pair (12,13) none of the randomly chosen 1kb regions correlate with a coefficient of >= 0.5. It is clear from these data that correlations between promoter regions, and to a smaller extent regions downstream of the gene, are distinctly different from correlating randomly selected 1kb regions. We also compared the motif vectors correlations (Pearson’s correlation) in the promoter regions of all genes between chr12-chr17 and chr18- chr19 with Spearman’s rank correlation. We observed that the trend was similar for both chromosome pair 12-17 and 18-19 with R^2^ values 0.83 and 0.79 respectively for 1kb 5-mer motif vector comparisons (supplementary figure 9 and supplementary data of Spearman’s vs Pearson’s correlations).

**Table 1a:**
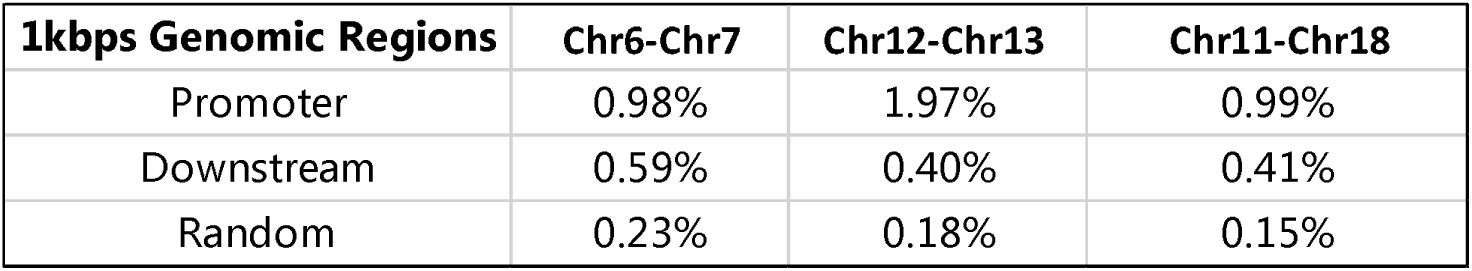
Comparison of gene-pair correlation above 0.5 in promoter and downstream vs random bins.

| **1kbps Genomic Regions** | **Chr6-Chr7** | **Chr12-Chr13** | **Chr11-Chr18** |
| --- | --- | --- | --- |
| Promoter | 0.98% | 1.97% | 0.99% |
| Downstream | 0.59% | 0.40% | 0.41% |
| Random | 0.23% | 0.18% | 0.15% |

**Table 1b:** Distribution of genes with overlapping coordinates in different chromosomes.

| **Chromosome** | **Total Genes** | **% Overlap** |
| --- | --- | --- |
| Chr 6 | 3086 | 22.65% |
| Chr 7 | 2891 | 21.90% |
| Chr 11 | 2995 | 20.50% |
| Chr 12 | 2575 | 24.82% |
| Chr13 | 1768 | 18.27% |
| Chr 18 | 1022 | 22.20% |

We also identified that the high abundance (OE >=5) of motifs of size 5 and 6 in the promoter regions holds true for a majority (∼ 50%) of the genes (supplementary table 3). The large number of genes whose promoters have a high abundance of motifs is suggestive that these motifs contribute to the recognition by some cognate protein regulators. On inspecting the motif distribution in promoter regions of 1kbps, we observed that on average ∼332 motifs out of 512 5-mer motif pairs and ∼1962 out of 2080 6-mer motif pairs have OE_1kbps_5/6-mer_ >= 5 in at least one promoter region. These motifs having OE_1kbps_5/6-mer_ >=5 across different promoter-proximal control regions contribute significantly to the correlations. We have also identified seven 5-mers and sixteen 6-mers that have OE_1kbps_5/6-mer_ >= 10 in at least 1% of the total genes in each chromosome (supplementary figure 10). These motifs show OE_1kbps_5/6-mer_ >= 10 consistently across all the chromosomes. The seven 5-mer conserved motifs are a subset of the sixteen 6-mer motifs thus suggesting the conservation of the base specific recognition by the regulatory partners. The seven 5-meric motifs (CGGGG, GCGCG, GCGGG, GGCGG, GGGGC, GGGGG, GGAGG) are all GC motifs (with the exception of GGAGG). Even though ∼165 and ∼1178 different combinations of 5-mer and 6-mer motifs show a high abundance (OE_1kbps_5/6-mer_ >= 10), only seven and sixteen of them are conserved across promoter proximal control regions. This implies that for differential regulation, different combinations of these high abundance motifs contribute to the high correlations.

### 4. Probing the significance of high motif vector correlations

#### 4.1 The relevance of high correlations

We looked at 3 genes whose promoter motif vectors had high correlations (> 0.9) to one another. These genes – LPHN1(ADGRL1) (adhesion G protein-coupled receptor L1), CDK9 (cyclin dependant kinase 9) and TRIM8 (tripartite motif containing 8) were from chromosomes 19, 9 and 10 respectively. The correlations between the motif vectors of (LPHN1 and CDK9), (LPHN1 and TRIM8) and (CDK9 and TRIM8) were 0.905, 0.903 and 0.910 respectively (supplementary table 4). The reason for the high correlation between these motif vectors is because of a few motifs that are common to all three with high OE ratio values (supplementary table 5). We next employed NetworkAnalyst^36,57^ to determine if there were common transcription factors to these three genes (figure 4a). Among the many transcription factors associated with these genes, two – SP1 and TFAP2A were common to all three genes. We use JASPER^58–60^ to get the consensus sequence recognised by these transcription factors (supplementary table 5).

**Figure 4:**
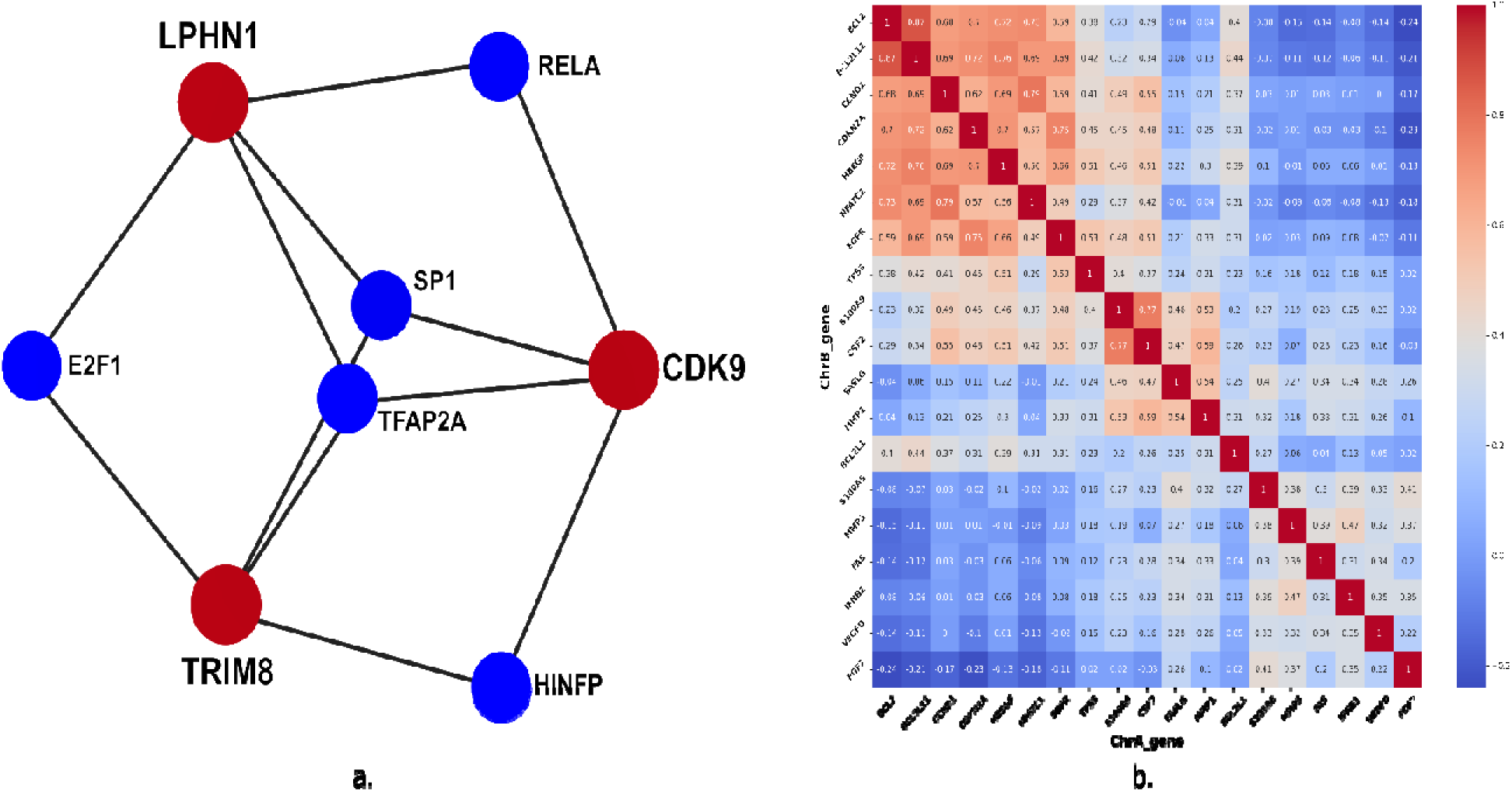
Panel a) is a graph representation of a gene regulatory network involving 3 genes LPHN1, CDK9 and TRIM8 (shown as red nodes). Each such red node is connected via edges to transcription factors controlling their expression (cyan nodes). Panel b) shows a gene-gene interaction heat map of OE_1kb_5-mer_ for 18 genes that are all regulated by the transcription factor Jun-Fos. The higher (or lower) the correlation the darker the shade of red (or bluer) is the grid.

The TF consensus for SP1 and TFAP2A from JASPAR can be interpreted as NNGGNNN[G/T][G/C/A][T/A] and GCCNNN[G/A][G/A/T][G/C] respectively. For SP1, a subset of three 5-mer motifs that can be derived from the consensus (GGCGG, CGGGG, GCCGG) have OE values above 10. Similarly, for TFAP2A, seven 5-mer motifs (GCGGC, GGGCG, GCCCG, GGGCC, GGCCG, GCCGG, GGGGC) derived from the consensus have OE values above 10 in all three gene promoters (supplementary table 5). It is important to note here that there were other 5-mers apart from those matching with the consensus that also had high OE values (supplementary table 5). From these data, it is clear that both the common transcription factors have an abundant number of binding sites on the promoters of all three genes.

An inspection of the tissues in which these three genes and their common two transcription factors are expressed reveals an interesting fact. The genes are almost ubiquitously expressed and the same is true for the transcription factor SP1(https://www.ncbi.nlm.nih.gov/gene/6667). TFAP2A is however expressed in fewer tissue types (https://www.ncbi.nlm.nih.gov/gene/7020) with a high expression in skin, placenta and salivary gland. For instance, LPHN1, CDK9 and TRIM8 are all implicated in functions related to fat ^61–65^. TFAP2A is expressed at low levels in adipose tissue. We conjecture that in fat, SP1 is the common regulator of all three genes. In other tissues, both SP1 and TFSP2A could assist in transcription, perhaps redundantly.

We also examined the relative abundances of motifs in the promoters for TFs that were common to two of the three proteins. We looked for motifs that were abundant in the promoters of two of the genes while being poorly represented in the other (supplementary table 5). The TFs – E2F1, HINFP and RELA are the TFs that are common between (LPNH1 and TRIM8), (CDK9 and LPNH1) and (CDK9 and TRIM8) respectively. The JASPAR consensus motifs for these TFs can be interpreted as TTT[G/C][G/C]CG[C/G], [A/G]C/G]GTCCGC and [G/C/T][G/T]G[A/G]NTTTCC respectively. Though there are TF consensus derived motifs that have higher OE ratio values in the promoters of the two genes sharing the same TF than the one not in common, the absolute values of the abundances are somewhat low. We noticed this for all three TFs E2F1, HINFP and RELA where 3, 5 and 1 motifs showed such relative abundances respectively. This may be indicative of the fact that differential regulation may not always involve TFs alone. As pointed out in the case of SP1 and TFAP2A, there are other motifs that are common to the promoters and present in high abundance but not linked to the TF. These are likely to be motifs that are recognized by other regulatory elements.

The illustration (figure 4) with the three genes and two common transcription factors exemplifies the mechanism of differential control of gene regulation. To ensure that regulators, such as transcription factors, bind to the promoter region there is an abundance of such binding sites on the promoter. In the 1kbps promoter regions of LPHN1, CDK9 and TRIM8 there are (22, 22), (31, 16), and (16, 20) binding sites for (SP1, TFAP2A) respectively. In general, the larger the number of such binding sites in the promoter region, the greater the probability of binding of the regulatory element. The redundancy in binding sites also ensures robustness in the case of mutations. The composite set of motifs in promoter regions is such that their extent of correlation to other promoter regions varies. On one end of the spectrum, there are high correlations, where the same set of motifs are repeated several times in a pair (supplementary table 3 and supplementary figure 10). Then there are correlations that are just high enough to be above a threshold. In such cases, it is likely that a pair shares a few common motifs of high OE ratios while there are others with high OE ratios in one promoter and not the other (supplementary table 5). A single promoter could hence have motifs in common with several other promoters which in turn may not be correlated to one another. This would form the basis of differential regulation ^66^.

#### 4.2 Constructing a gene regulatory network

Following up on the theme of how differential regulation can be affected in cells, we looked at the promoter regions of all genes that were known to be transcribed by the same transcription factor(s) – Jun/Fos^67,68^. We selected a set of 19 genes whose transcription is controlled by the transcription factors Jun/Fos ^69^. The genes were identified in mouse while our analysis considered the human homologous of the mouse genes.

An all-vs-all correlation was done using the promoter motif vectors over this gene set (figure 4b). Some of the genes are more closely correlated with a few of the other genes and the pattern of clusters that emerge has 3 major groups of 10, 4 and 6 genes. The genes were clustered into 3 groups based on the genes which have higher correlations within themselves compared to others in the list of genes. Within each cluster, the genes are better correlated to one another than they are to the genes in other clusters. There is a nuanced patterning of the high-scoring common motifs among these gene clusters that lead to higher intra-cluster correlation values. An interesting observation here is that the Fos/Jun binding consensus motif TGACTCA is not among the motifs with high OE ratio values. The motifs with high OE here that are in common to several genes (such as motif CGCGG present in genes BCL2, BCL2L11, HBEGF and EGFR) could be binding sites of some gene regulators (identity unknown) (supplementary data of AP1 network). This example shows how one could construct a gene regulatory network by clustering together genes whose promoters share the same motifs.

#### 4.3 Spatially proximal genes have strong motif vector correlations

We rationalised that genes whose promoters are correlated are likely to be coregulated. Our next investigation was to check how often correlation/coregulation implied colocalization and whether we could predict it. Experimentally, colocalization is inferred using chemical cross linking such as in Hi-C ^9,16^. Data from Hi-C experiments are at the resolution of a few kbps or even tens of kbps. Our computations however are at a higher resolution as our inferences are drawn from 5/6 base motifs. More precisely, we are interested in the abundances of all 5/6-mer motifs in a given sequence stretch, variously considered as 1kbps, 2kbps and 6kbps immediately upstream of genes. We also looked at promoter capture Hi-C data (pcHi-C), which captures distal promoter interacting regions for all promoters^70^. We looked at the correlation of genes that overlap with these regions detected by Hi-C experiments and how they are correlated at different resolutions. In this study, we used two datasets: 1. Hi-C data of long-range chromatin contacts in HCT116 colon cancer cells ^37^(accessible under GEO accession number GSE18215) and 2. pcHi-C data, which looks at novel gene contacts associated with autoimmune risk loci^38^ (accessible under GEO accession number: GSM1704495).

Three important caveats apply to our analysis: 1). Hi-C records cross links across the genome without any prejudice to any region(s). Our data, in this study, is restricted to correlations in gene promoter regions only. This implies that we would only be able to compare our correlation data to a subset of Hi-C data that cross link promoter regions of genes. We make a higher number of predictions as we consider all possible gene pair correlations between the two chromosomes. 2). The resolution of data for the both the Hi-C and the pcHi-C experiments are in the order of ∼10 kbps, whereas our predictions, that examine 5/6 bps, are of higher resolution. 3). Different Hi-C experiments use different genome reference assemblies to establish the genomic coordinates between two chromosomes that are in proximity. The gene annotations vary with each reference assembly (supplementary table 6). We have used all the gene annotations according to the latest T2T reference assembly. Thus, it is likely that we are missing out on some gene correlations, because of discrepancies in the annotations across different reference assemblies.

Here we have compared our motif vector correlations to Hi-C data for a couple of chromosomes – 18 and 19 that have 1022 (spread over ∼80.5mbps) and 2531 genes (spread over ∼61.7mbps) respectively. These two chromosomes were chosen because they localize differently in the nucleus – 18 is peripheral while 19 is mostly internalised, ^71,72^ and have contrasting gene densities^73^ (Supplementary figure 3). The Hi-C data provides the contact links for different parts of the two chromosomes. We obtained the coordinates of the contacts between the two chromosomes and obtained the genes whose coordinates overlap with the Hi-C contacts. The extraction of the gene information was done using the respective reference assembly that was used to generate the Hi-C data.

According to the 10kb bin long range chromatin contact Hi-C data, there are 1,327,617, 699,290 and 23,688 Hi-C contacts intra-chromosome-18, intra-chromosome-19 and inter-18- 19 respectively according to the GRCh38 reference assembly. Of these, only 12,790, 115,595 and 926 are contacts overlap with regions having genes in intra-18, intra-19 and inter-18-19 respectively. Overlap with gene pairs is established when both Hi-C contact bins have a gene (in whole or part). Expectedly, the number of inter-18-19 contacts is small in comparison to the intra-chromosome contacts given their different localizations. The aim here was to determine if the gene pairs whose promoter motif vector are correlated are also physically proximal. Of the overlaps determined, only 12,213, 111,730 and 890 gene-pairs find a match with the current T2T gene annotation list which is our reference for the motif vector correlations (supplementary data of Hi-C validation).

We identify the genes having motif vector correlation scores >= 0.5 from the inter and intra chr18, chr19 Hi-C contacts. We see enrichment in the motif vector correlations in the Hi-C contacts for all the inter and intra-Hi-C contacts at all resolutions of motif vector correlations at different sizes (Table 2). We also verified if the high correlations between the gene promoter are a result of sequential proximity between the genes/promoters. As a control, we plotted the distance between the genes against the correlation of their promoter regions. We observe that high correlations between gene promoter regions are not dependent on the sequential proximity of the genes (supplementary figure 11). It is to be noted that this comparison was done only across intra-gene correlations as we cannot determine the sequential proximity of inter-chromosome correlations.

**Table 2:** Comparison of gene-pair correlation inter and intra chromosome 18 and chromosome 19 in Hi-C contacts vs all.

| **Chr-Pair** | Gene pairs with motif vector<br>correlation $\geq 0.5$ | | Gene pairs in **Hi-C** with motif vector<br>correlation $\geq 0.5$ | |
| --- | --- | --- | --- | --- |
|  | **1kb 5-mer** | **1kb 6-mer** | **1kb 5-mer** | **1kb 6-mer** |
| **Chr 18 - Chr 19** | 9.92% | 0.47% | 14.38% | 1.01% |
| **Chr 18 – Chr 18** | 9.96% | 1.18% | 18.84% | 7.72% |
| **Chr 19 – Chr 19** | 15.10% | 0.90% | 22.18% | 3.74% |

We repeated this analysis using pcHi-C data using chromosome pairs 18 and 19 and obtained 3655 contacts. Of these, 123 gene coordinates overlap with the pcHi-C contacts according to the GRCh37 reference assembly (supplementary table 7 and supplementary data of Hi-C validations). 118 of the 123 gene-pairs match with our correlations using current T2T reference assembly gene annotations (table 3 and supplementary tables 6).

**Table 3:**
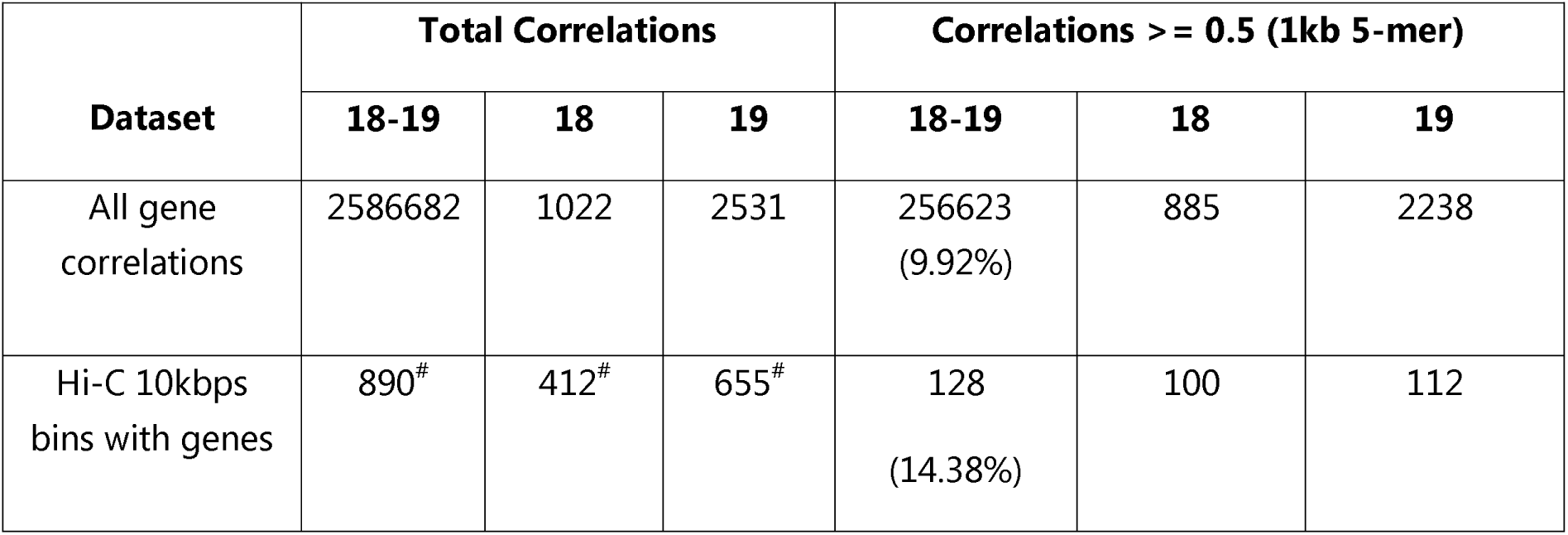
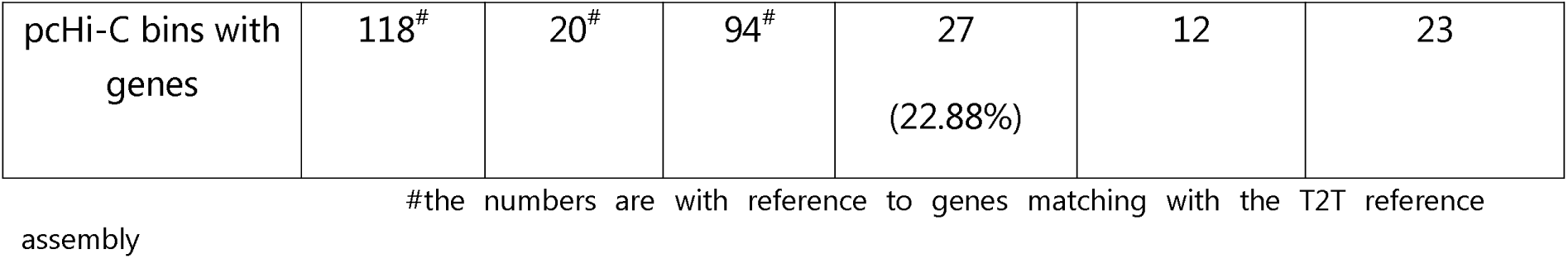
Distribution of gene pairs contributing to high correlation in Hi-C and pcHi-C.

Of all the 2,586,682 possible gene pairs of Chromosomes 18 and 19, 256,623 (9.92% of 1kb_5-mer) have motif vector correlations of >= 0.5 (supplementary table 7). 128 (14.32%) gene pairs with Hi-C contacts have a motif vector correlation >= 0.5, an enrichment of ∼5%. This number is 27 (∼23%%) for pcHi-C. Interestingly there is no overlap between the Hi-C and the pcHi-C data (supplementary figure 12). Cumulatively, the motif vector correlations of >= 0.5 identify 155 pairs of genes that are experimentally linked to one another, which is an enrichment of 37%. Given that the two different types of Hi-C have no overlaps, it is possible that many more of the gene pairs identified by our correlations could be spatially proximal under certain conditions.

Topologically associated domains (TADs) are well defined chromosome segments that make more frequent genomic contacts to one another and are hence purportedly in spatial proximity^74^. It has been established that binding sites for the protein CTCF, characterised by the motif CCCTC, mark the TAD boundaries^75,76^. The CCCTC motif is among the top 50 most abundantly present motifs in the human genome (43^rd^ on a list of 512 with an average OE_1kbps_5me_ value of 1.85 (supplementary data of OE_whole_ rations). On closer inspection, we observed that 349 (39.2%) and 67 (56.8%) out of the 890 and 118 gene pairs in pcHi-C and Hi-C respectively show an OE_1kbps_5-mer_ of the CTCF motif greater than 3 in both of the genes in contact (supplementary data of Hi-C validation). This number is enriched when we look at gene pairs with correlation of >= 0.5. We observed that 93 (72.6%) and 23 (85.2%) out of the 128 and 27 gene pairs with correlation >= 0.5 in pcHi-C and Hi-C respectively show an OE_1kbps_5-mer_ of the CTCF motif greater than 3 in both of the genes. Interestingly, the OE_1kbps_5-mer_ for CCCTC in the promoter regions of gene pairs that have a motif vector correlation of >= 0.5 and are proximal to Hi-C or the pcHi-C contacts in chromosomes 18 and 19 has a maximum value of ∼20, with a mean of ∼3.5 to 4.5. (Supplementary figure 13 and supplementary data of Hi-C validation). In contrast, the OE_1kbps_5-mer_ for the other motifs in the genes in proximity, do not exceed a maximum of 2.5, with their means ranging from ∼1 to 1.5. Thus, the CTCF motif is one of the major contributors to the high correlation (correlation coefficient >= 0.5) for gene pairs in proximity.

The motif vector correlations are high among gene pairs that also have Hi-C contacts. This opens up the possibility that Hi-C contacts could be predicted from gene correlations. There is no overlap between the 123 and 926 gene pairs identified by Hi-C and pcHi-C respectively. Though these data involve 15 and 33 genes from chromosomes 18 and 19 respectively, they are always matched with different partner genes in the two sets of data (supplementary figure 12).

## DISCUSSIONS

In this study, we have attempted to read parts of the genome just as they would be perceived by the proteins that interact with them, i.e., 4 to 6 base pairs at a time. These are data we have obtained by observing DNA-protein complexes. For our analysis, we simplistically and separately contend that DNA binding proteins recognize their cognate sites on genomes by making interaction with either 5 or 6 bases per domain. Some databases list consensus binding sites that are longer – but these are usually in cases where the DNA binding domains dimerise.

How are 5/6 mers of DNA distributed in the genome? There are a total of 512 and 2080 unique 5-mers and 6-mers respectively that were obtained by averaging the values of motifs and their corresponding reverse complements. We have shown that the distribution of these 5/6-mers is non-random. In real genomes, there are regions of the genome where certain motifs are over or under-represented. We have quantified this using the OE ratios. The non-randomness of the motif distributions happens at the scale of the whole chromosome or even at the level of smaller segments of the genome such as 100kbps stretches or even around centromeric regions across all chromosomes. Using the simple metric of OE_centromere_ ratios, we establish a unique pattern of motif distributions conserved across the different acrocentric chromosomes (13,14,15,21 and 22) The centromeres of these chromosomes are known to have segmental duplications and our analysis using only the motif distributions in these regions also identifies similarities.

Having established that the motif distributions follow non-random patterns we next investigated its possible relevance. For this, we first correlated (Pearson correlation) the motif distributions in promoter regions of genes. To do this we constructed motif vectors of the 5/6-mer motifs where the coefficients of the vectors are the OE ratios of the motifs. When the Having established that the motif distributions follow non-random patterns we next investigated its possible relevance. For this, we first correlated (Pearson correlation) the motif distributions in promoter regions of genes. To do this we constructed motif vectors of the 5/6-mer motifs where the coefficients of the vectors are the OE ratios of the motifs. We observed that certain 5- and 6-mer motifs have high abundance across the majority of the promoter regions of genes (∼50%). We identified seven 5-mers and sixteen 6-mers that are conserved across the promoter regions in different chromosomes. These motifs of high abundance suggest identification by some cognate protein regulators. When the correlation values of all genes are taken together chromosome-wise, we observed that there were some chromosome pairs with very strong correlations (>= 0.9). We have highlighted two such clusters of high correlations – between chromosomes (13,14,15,21 and 22) and (4,10). In both these cases there are documented instances of chromosome translocations amongst cluster members. The (13,14,15,21,22) clusters are the same acrocentric chromosomes discussed above. The mix and match of chromosomal segments between these genes is termed Robertsonian translocations and is implicated in genetic diseases such as Patau syndrome^77^ and Down syndrome^78^. Translocations between chromosomes 4 and 10 are associated with certain types of leukaemia^54,55^ and Wolf-Hirschhorn syndrome^56^. Our somewhat simplistic motif vector correlations also pick up on these phenomena. Translocations may only be possible when there is a high degree of sequence similarity/identity between the regions of translocation. However, not all regions sharing high similarity/identity may be susceptible to translocations as it may require the presence of certain motif(s). What motif(s) and their distributions can only be gauged with a focused analysis of the translocating segments. The high abundances of promoter correlations are also an attribute of the tandem repeats in the promoter regions of the rRNA genes across these chromosomes. The patterns of high correlation across gene promoters are also distinct from that of downstream regions of genes or random regions in the chromosomes.

From the level of chromosomes, we went down to the level of individual genes, more precisely to the promoter region of genes. We showed that when two genes have high correlations between their promoter vector motifs there is usually a functional implication. For instance, the genes LPNH1 from chromosome 19, CDK9 from chromosome 9 and TRIM8 from chromosome 10 are all strongly correlated to one another (correlation coefficient > 0.9). The reason for the high correlations there is a common set of motifs that are over/under-represented in all their promoter regions. Examining the over-abundant motifs, we found that the motifs GGCGG and CCGGG were among the ones with the highest OE ratios in all their promoters and these were the binding sites of transcription factors SP1 and TFAP2A. These transcription factors are differentially expressed in different tissue types, for instance, TFAP2A is not found in fat while SP2 is. This, we speculate is the basis of differential gene regulation in different cell/tissue types. Further, we could construct an entire gene regulatory network by associating gene promoter regions with significant correlations. While we are constructing this network, it is too vast and somewhat beyond the scope of the current investigation to be reported here.

Having introduced the possibility of obtaining gene regulatory networks, we illustrate this with an example of promoter correlations between all co-regulated genes, i.e., genes transcribed by a common transcription factor, Jun/Fos. Here, we found some gene clusters to be strongly correlated amongst cluster members but not necessarily with members of other clusters. The reason for this is again the fact that the correlations are strong because of a common set of motifs having similar abundances (OE ratios) across promoter regions. These motifs are then of functional importance as established in the case of LPHN1, CDK9 and TRIM8. In this illustrative network, however, the common high abundance motifs are not transcription binding sites (even though some of the pairs within the network share common transcription factors other than Jun/Fos). The Jun/Fos binding motifs themselves are not among the motifs with the highest ratios. We speculate that the over-abundant common set of motifs could be binding sites for common regulatory elements, perhaps hitherto undiscovered. Since the high-abundance motifs in promotors within a single cluster are not identical between cluster members, it opens up the possibility of differential regulation.

Our final investigation was to examine whether high gene promoter correlation also implied physical proximity. Here the results are interesting and suggestive of the predictive power of these motif vector correlations. On correlating the results to Hi-C data that show physical proximity between genic regions using cross-linking, we find that in an enrichment of the high motif vector correlations (coefficients >= 0.5) overlap with Hi-C data. This enrichment was consistent when the analysis was repeated with pcHi-C. While processing this result, we should bear in mind that the number of gene-gene promoter correlations with coefficients >0.5 is at least an order of magnitude greater than the amount of Hi-C data available to us. While it is unlikely that all strongly correlated gene-gene promoter regions imply physical proximity, it is possible that many of them do, even though there is no Hi-C data to support the claim. The reason for this could be that Hi-C data is typically of a resolution of 10kbps (in many cases >10kbps), while we are examining the genome at a higher resolution (length scale of 5/6 base pairs). The promoter regions of the genes in Hi-C contacts also showed a high abundance of the CTCF binding motifs that regulates the TAD organisation. One significant observation we made was the lack of overlap between the different Hi-C data. The gene pairs present in the bins with the Hi-C contacts do not overlap with the pcHi-C data. This can be an attribute of the difference in the reference assembly used to generate the Hi- C and also the possible different crosslinks in different cell types. Our observations on high correlations are made with the latest reference assembly, and are independent of cell type. One other reason why some of our high correlations are not validated by Hi-C data could be that even though the genes are in close proximity, they may not have been close enough to be detected by Hi-C, but could be close enough to be co-regulated. We are also aware that spatial proximity may not always follow from strong motif vector correlations even though the genes are co-regulated. For instance, transcription factors that have peripheral locations on the nucleus could co-regulate two (or more) genes that are spatially distant but are also somewhat close to the nuclear outer periphery. These results open up the possibility of predicting, at a high resolution, 3D proximities within genomes. Potentially, this could give us a more nuanced map of the genomic arrangement within the nucleus and how it changes from one cell type to another or even during different cell phases.

Typically, analysis of genes/promoters/genomes etc. involves sequence alignments. While the benefits of alignments are undeniable, this study shows that it is insightful and significant to read the genome 5/6 bases at a time, just as proteins that bind DNA do. From a rather simplistic consideration of the distribution of 5/6-mers in the chromosome or smaller stretches of the genomes (such as promoter regions), we can make fundamental connections between genes and chromosomes. For instance, given that we can detect patterns where translocations are likely to occur, predict the proximity of correlated (co-regulated) genes, and give ourselves the ability to construct regulatory networks, we see great benefit in examining motif abundances in the genome.

## DATA AVAILABILITY

http://sites.iiserpune.ac.in/~madhusudhan/DNA_motifs/Supplementary_data_NAR_06_09_24. zip

## AUTHOR CONTRIBUTIONS

Idea conception: MSM with help from AC. Data generation and analysis: AC with help from SC. Manuscript writing: AC and MSM

## ACKNOWLEDGEMENTS

We would like to acknowledge Prof. Chandra Verma, Prof. Kundan Sengupta, Prof. Richa Ricky, Prof. Leelavati Narlikar, S. Mukundan, Avadhoot Jadhav and other COSPI lab members for constructive criticisms. We also acknowledge the support and computational resources provided by the PARAM Brahma facility under the National Supercomputing Mission, Government of India at the Indian Institute of Science Education and Research (IISER) Pune.

## REFERENCES

1. Aganezov, S. et al. A complete reference genome improves analysis of human genetic variation. Science (80-.). 376, (2022).

2. Nurk, S. et al. The complete sequence of a human genome. Science (80-.). 376, 44–53 (2022).

3. Dunham, I. et al. An integrated encyclopedia of DNA elements in the human genome. Nature 489, 57–74 (2012).

4. Bejerano, G., Haussler, D. & Blanchette, M. Into the heart of darkness: large-scale clustering of human non-coding DNA. Bioinformatics 20 Suppl 1, (2004).

5. Guturu, H., Doxey, A. C., Wenger, A. M. & Bejerano, G. Structure-aided prediction of mammalian transcription factor complexes in conserved non-coding elements. Philos. Trans. R. Soc. Lond. B. Biol. Sci. 368, (2013).

6. Fedoroff, N. V. Transposable elements, epigenetics, and genome evolution. Science (80-.). 338, 758–767 (2012).

7. Vitsios, D., Dhindsa, R. S., Middleton, L., Gussow, A. B. & Petrovski, S. Prioritizing non-coding regions based on human genomic constraint and sequence context with deep learning. Nat. Commun. 2021 121 12, 1–14 (2021).

8. Sémon, M. & Duret, L. Evolutionary origin and maintenance of coexpressed gene clusters in mammals. Mol. Biol. Evol. 23, 1715–1723 (2006).

9. Schneider, V. A., et al. EvSchneider, V. A., Graves-Lindsay, T., Howe, K., Bouk, N., Chen, H. C., Kitts, P. A., Murphy, T. D., Pruitt, K. D., Thibaud-Nissen, F., Albracht, D., Fulton, R. S., Kremitzki, M., Magrini, V., Markovic, C., McGrath, S., Steinberg, K. M., Auger, K., Chow,. Genome Res. 27, 849–864 (2017).

10. Church, D. M. A next-generation human genome sequence. Science (80-.). 376, 34–35 (2022).

11. Cozzolino, F., Iacobucci, I., Monaco, V. & Monti, M. Protein-DNA/RNA Interactions: An Overview of Investigation Methods in the -Omics Era. J. Proteome Res. 20, 3018–3030 (2021).

12. Dekker, J. & Mirny, L. The 3D Genome as Moderator of Chromosomal Communication. Cell 164, 1110–1121 (2016).

13. Comfort, N. C. From controlling elements to transposons: Barbara McClintock and the Nobel Prize. Trends Genet. 17, 475–478 (2001).

14. Misteli, T. Higher-order genome organization in human disease. Cold Spring Harb. Perspect. Biol. 2, (2010).

15. de Wit, E. & de Laat, W. A decade of 3C technologies: insights into nuclear organization. Genes Dev. 26, 11–24 (2012).

16. Tjong, H. et al. Population-based 3D genome structure analysis reveals driving forces in spatial genome organization. Proc. Natl. Acad. Sci. U. S. A. 113, E1663–E1672 (2016).

17. Van Steensel, B. & Henikoff, S. Identification of in vivo DNA targets of chromatin proteins using tethered Dam methyltransferase. Nat. Biotechnol. 18, 424–428 (2000).

18. Guelen, L. et al. Domain organization of human chromosomes revealed by mapping of nuclear lamina interactions. Nature 453, 948–951 (2008).

19. Solomon, M. J., Larsen, P. L. & Varshavsky, A. Mapping protein-DNA interactions in vivo with formaldehyde: evidence that histone H4 is retained on a highly transcribed gene. Cell 53, 937–947 (1988).

20. Orlando, V. Mapping chromosomal proteins in vivo by formaldehyde-crosslinked-chromatin immunoprecipitation. Trends Biochem. Sci. 25, 99–104 (2000).

21. Buenrostro, J. D., Giresi, P. G., Zaba, L. C., Chang, H. Y. & Greenleaf, W. J. Transposition of native chromatin for fast and sensitive epigenomic profiling of open chromatin, DNA-binding proteins and nucleosome position. Nat. Methods 10, 1213–1218 (2013).

22. Henikoff, S., Henikoff, J. G., Kaya-Okur, H. S. & Ahmad, K. Efficient chromatin accessibility mapping in situ by nucleosome-tethered tagmentation. Elife 9, 1–19 (2020).

23. Nguyen, H. Q. et al. 3D mapping and accelerated super-resolution imaging of the human genome using in situ sequencing. Nat. Methods 2020 178 17, 822–832 (2020).

24. Nir, G. et al. Walking along chromosomes with super-resolution imaging, contact maps, and integrative modeling. PLOS Genet. 14, e1007872 (2018).

25. Boninsegna, L. et al. Integrative genome modeling platform reveals essentiality of rare contact events in 3D genome organizations. Nat. Methods 2022 198 19, 938–949 (2022).

26. Vinga, S. & Almeida, J. Alignment-free sequence comparison—a review. Bioinformatics 19, 513–523 (2003).

27. Reinert, G., Chew, D., Sun, F. & Waterman, M. S. Alignment-free sequence comparison (I): statistics and power. J. Comput. Biol. 16, 1615–1634 (2009).

28. Sarkar, B. K. et al. Determination of k-mer density in a DNA sequence and subsequent cluster formation algorithm based on the application of electronic filter. Sci. Reports 2021 111 11, 1–12 (2021).

29. Saw, A. K. et al. Alignment-free method for DNA sequence clustering using Fuzzy integral similarity. Sci. Rep. 9, (2019).

30. Nair, S. & Madhusudhan, M. S. JEDII: Juxtaposition Enabled DNA-binding Interface Identifier. bioRxiv 2022.05.19.492702 (2022) doi:10.1101/2022.05.19.492702.

31. Wingender, E. Compilation of transcription regulating proteins. Nucleic Acids Res. 16, 1879–1902 (1988).

32. Wingender, E., Schoeps, T., Haubrock, M., Krull, M. & Dönitz, J. TFClass: expanding the classification of human transcription factors to their mammalian orthologs. Nucleic Acids Res. 46, D343–D347 (2018).

33. Rohs, R. et al. The role of DNA shape in protein–DNA recognition. Nat. 2009 4617268 461, 1248–1253 (2009).

34. Chiu, T. P., Xin, B., Markarian, N., Wang, Y. & Rohs, R. TFBSshape: an expanded motif database for DNA shape features of transcription factor binding sites. Nucleic Acids Res. 48, D246–D255 (2020).

35. Zhou, G. et al. NetworkAnalyst 3.0: a visual analytics platform for comprehensive gene expression profiling and meta-analysis. Nucleic Acids Res. 47, W234–W241 (2019).

36. Xia, J., Gill, E. E. & Hancock, R. E. W. NetworkAnalyst for statistical, visual and network-based meta-analysis of gene expression data. Nat. Protoc. 10, 823–844 (2015).

37. Spracklin, G. et al. Diverse silent chromatin states modulate genome compartmentalization and loop extrusion barriers. Nat. Struct. Mol. Biol. 2022 301 30, 38–51 (2022).

38. Martin, P. et al. Capture Hi-C reveals novel candidate genes and complex long-range interactions with related autoimmune risk loci. Nat. Commun. 2015 61 6, 1–7 (2015).

39. Homo sapiens genome assembly T2T-CHM13v2.0 - NCBI - NLM. https://www.ncbi.nlm.nih.gov/datasets/genome/GCF_009914755.1/.

40. NCBI Genome Data Viewer. https://www.ncbi.nlm.nih.gov/genome/gdv/.

41. Nassar, L. R. et al. The UCSC Genome Browser database: 2023 update. Nucleic Acids Res. 51, D1188–D1195 (2023).

42. Gonzalez, D. et al. ZNF143 protein is an important regulator of the myeloid transcription factor C/EBPα. J. Biol. Chem. 292, 18924 (2017).

43. Humphray, S. J. et al. DNA sequence and analysis of human chromosome 9. Nature 429, 369–374 (2004).

44. Sinclair, D. A. R., Schulze, S., Silva, E., Fitzpatrick, K. A. & Honda, B. M. Essential genes in autosomal heterochromatin of Drosophila melanogaster. Genetica 109, 9–18 (2000).

45. Eberl, D. F., Duyf, B. J. & Hilliker, A. J. The role of heterochromatin in the expression of a heterochromatic gene, the rolled locus of Drosophila melanogaster. Genetics 134, 277–292 (1993).

46. Rhie, A. et al. The complete sequence of a human Y chromosome. Nature 621, 344 (2023).

47. Stadhouders, R. et al. Transcription factors orchestrate dynamic interplay between genome topology and gene regulation during cell reprogramming. Nat. Genet. 50, 238 (2018).

48. Groves, A. K. et al. Differential regulation of transcription factor gene expression and phenotypic markers in developing sympathetic neurons. Development 121, 887–901 (1995).

49. Hartley, G. & O’neill, R. J. Centromere Repeats: Hidden Gems of the Genome. Genes (Basel). 10, (2019).

50. Vollger, M. R. et al. Segmental duplications and their variation in a complete human genome. Science (80-.). 376, (2022).

51. Hurst, L. D., Sachenkova, O., Daub, C., Forrest, A. R. R. & Huminiecki, L. A simple metric of promoter architecture robustly predicts expression breadth of human genes suggesting that most transcription factors are positive regulators. Genome Biol. 15, 1– 26 (2014).

52. Saeed, S., Hassan, J., Javed, S. M., Shan, S. & Naz, M. A Familial Case of Robertsonian Translocation 13;14: Case Report. Cureus 14, (2022).

53. Spinner, N. B., Conlin, L. K., Mulchandani, S. & Emanuel, B. S. Deletions and Other Structural Abnormalities of the Autosomes. Emery Rimoin’s Princ. Pract. Med. Genet. 1– 37 (2013) doi:10.1016/B978-0-12-383834-6.00051-3.

54. Harris, M. et al. Trisomy of Leukemic Cell Chromosomes 4 and 10 Identifies Children With B-Progenitor Cell Acute Lymphoblastic Leukemia With a Very Low Risk of Treatment Failure: A Pediatric Oncology Group Study. Blood 79, 3316–3324 (1992).

55. Wong, K. F. & So, C. C. Acute myeloid leukemia with concomitant trisomies 4 and 10: a distinctive form of myeloid leukemia? Cancer Genet. Cytogenet. 127, 74–76 (2001).

56. Goodship, J. et al. A submicroscopic translocation, t(4;10), responsible for recurrent Wolf-Hirschhorn syndrome identified by allele loss and fluorescent in situ hybridisation. J. Med. Genet. 29, 451 (1992).

57. Xia, J., Benner, M. J. & Hancock, R. E. W. NetworkAnalyst - Integrative approaches for protein-protein interaction network analysis and visual exploration. Nucleic Acids Res. 42, (2014).

58. A., M., et al. JASPAR 2014: an extensively expanded and updated open-access database of transcription factor binding profiles. Nucleic Acids Res. 42, D142–D147.

59. Mathelier, A., et al. JASPAR 2014: An extensively expanded and updated open-access database of transcription factor binding profiles. Nucleic Acids Res. 42, (2014).

60. Castro-Mondragon, J. A., et al. JASPAR 2022: the 9th release of the open-access database of transcription factor binding profiles. Nucleic Acids Res. 50, D165–D173 (2022).

61. Martinez, A. F., Muenke, M. & Arcos-Burgos, M. From the Black Widow Spider to Human Behavior: Latrophilins, a Relatively Unknown Class of G Protein-Coupled Receptors, Are Implicated in Psychiatric Disorders. Am. J. Med. Genet. B. Neuropsychiatr. Genet. 0, 1 (2011).

62. Itkonen, H. M., Poulose, N., Walker, S. & Mills, I. G. CDK9 Inhibition Induces a Metabolic Switch that Renders Prostate Cancer Cells Dependent on Fatty Acid Oxidation. Neoplasia 21, 713–720 (2019).

63. Yan, F. J. et al. The E3 ligase tripartite motif 8 targets TAK1 to promote insulin resistance and steatohepatitis. Hepatology 65, 1492–1511 (2017).

64. Sayers, E. W. et al. Database resources of the national center for biotechnology information. Nucleic Acids Res. 50, D20–D26 (2022).

65. Fagerberg, L. et al. Analysis of the human tissue-specific expression by genome-wide integration of transcriptomics and antibody-based proteomics. Mol. Cell. Proteomics 13, 397–406 (2014).

66. Shabalina, S. A., Spiridonov, A. N., Spiridonov, N. A. & Koonin, E. V. Connections between Alternative Transcription and Alternative Splicing in Mammals. Genome Biol. Evol. 2, 791 (2010).

67. Shaulian, E. & Karin, M. AP-1 as a regulator of cell life and death. Nat Cell Biol 4, E131– E136 (2002).

68. Zenz, R. & Wagner, E. F. Jun signalling in the epidermis: from developmental defects to psoriasis and skin tumors. Int J Biochem Cell Biol 38, 1043–1049 (2006).

69. Zenz, R. et al. Activator protein 1 (Fos/Jun) functions in inflammatory bone and skin disease. Arthritis Res. Ther. 10, 1–10 (2008).

70. Schoenfelder, S., Javierre, B. M., Furlan-Magaril, M., Wingett, S. W. & Fraser, P. Promoter Capture Hi-C: High-resolution, Genome-wide Profiling of Promoter Interactions. J. Vis. Exp. 2018, 57320 (2018).

71. Croft, J. A. et al. Differences in the Localization and Morphology of Chromosomes in the Human Nucleus. J. Cell Biol. 145, 1119 (1999).

72. Boyle, S. et al. The spatial organization of human chromosomes within the nuclei of normal and emerin-mutant cells. Hum. Mol. Genet. 10, 211–220 (2001).

73. Vitalini, M. W., Dialynas, G., Wallrath, L. L., Mackey, S. R. & Stainbrook, S. C. Nucleus | Nuclear Organization, Chromatin Structure, and Gene Silencing. Encycl. Biol. Chem. Third Ed. 5, 393–397 (2021).

74. Sefer, E. A comparison of topologically associating domain callers over mammals at high resolution. BMC Bioinforma. 2022 231 23, 1–39 (2022).

75. Nanni, L., Ceri, S. & Logie, C. Spatial patterns of CTCF sites define the anatomy of TADs and their boundaries. Genome Biol. 21, 1–25 (2020).

76. Liu, Y. & Dekker, J. CTCF–CTCF loops and intra-TAD interactions show differential dependence on cohesin ring integrity. Nat. Cell Biol. 2022 2410 24, 1516–1527 (2022).

77. Kuznetsova, M. A., Zaytseva, G. V., Zryachkin, N. I., Makarova, O. A. & Khmilevskaya, S. A. Patau Syndrome. Vopr. Prakt. Pediatr. 10, 90–93 (2023).

78. Chung, W. K. Genetics. Fetal and Neonatal Secrets 266–289 (2014) doi:10.1016/B978-0-323-09139-8.00011-0.

