## supplementary for "Motif distribution in genomes gives insights into gene clustering and co-regulation"

*Supplementary Table1: Centromere boundaries for $OE_{centromere}$ calculation. Column 1 represents the gene name. Columns 2 and 3 represent the start-stop boundaries for each chromosome respectively. Columns 4 and 5 show the chromosome-wise centromere size in bps and Mbps respectively.*

| Chromosome | Start | Stop | Size(bps) | Size/Mbps |
|---|---|---|---|---|
| chr1 | 121973252 | 125421848 | 3448596 | 3.45 |
| chr2 | 91879019 | 96296965 | 4417946 | 4.42 |
| chr3 | 87983662 | 93990946 | 6007284 | 6.01 |
| chr4 | 48247029 | 51956625 | 3709596 | 3.71 |
| chr5 | 46259962 | 51541276 | 5281314 | 5.28 |
| chr6 | 58631981 | 62603545 | 3971564 | 3.97 |
| chr7 | 58365242 | 62251569 | 3886327 | 3.89 |
| chr8 | 43210183 | 47354764 | 4144581 | 4.14 |
| chr9 | 42215186 | 45645422 | 3430236 | 3.43 |
| chr10 | 38050203 | 41649353 | 3599150 | 3.6 |
| chr11 | 51079005 | 55858265 | 4779260 | 4.78 |
| chr12 | 33334329 | 37896869 | 4562540 | 4.56 |
| chr13 | 16502400 | 18949340 | 2446940 | 2.45 |
| chr14 | 16100180 | 18266678 | 2166498 | 2.17 |
| chr15 | 17458137 | 20558059 | 3099922 | 3.1 |
| chr16 | 35322745 | 38439712 | 3116967 | 3.12 |
| chr17 | 22694097 | 27672900 | 4978803 | 4.98 |
| chr18 | 15431301 | 21560433 | 6129132 | 6.13 |
| chr19 | 24221370 | 28153079 | 3931709 | 3.93 |
| chr20 | 25773360 | 30501863 | 4728503 | 4.73 |
| chr21 | 10901609 | 13034530 | 2132921 | 2.13 |
| chr22 | 13679758 | 17432904 | 3753146 | 3.75 |
| chrX | 58278961 | 63921024 | 5642063 | 5.64 |
| chrY | 10300948 | 10664448 | 363500 | 0.36 |

*Supplementary Table 2a: Chromosome-wise composition of rRNA genes*

| Chromosome | rRNA genes |
|---|---|
| 13 | 12.8% |
| 14 | 12.2% |
| 15 | 7.1% |
| 21 | 15.4% |
| 22 | 4.4% |

*Supplementary Table 2b: Proportion of rRNA genes contributing to high correlations (>=0.9)*

| Chromosome | 14 | 15 | 21 | 22 |
|---|---|---|---|---|
| 13 | 82.70% | 91.50% | 83.7% | 87.4% |
| 14 | | 81.50% | 81.5% | 80.9% |
| 15 | | | 86.10% | 87.5% |
| 21 | | | | 82.3% |

*Supplementary Table 3: Count of gene promoter proximal control elements having motifs with high $OE_{nkbps\_kmer}$. Column 1 represents the resolution of promoter and motif where n is promoter size and k is motif size. Columns 2,3,4 shows the result for OE cutoffs 5, 8 and 10 respectively.*

| **Resolution** | **OE >= 5** | **OE >= 8** | **OE >= 10** |
|---|---|---|---|
| 1kb_5-mer | 51,103 | 32,235 | 22,871 |
| 1kb_6-mer | 57,660 | 57,534 | 55,404 |
| 2kb_5-mer | 44,165 | 21,983 | 13,370 |
| 2kb_6-mer | 57,659 | 54,485 | 48,882 |
| 6kb_5-mer | 33,453 | 11,412 | 6,142 |
| 6kb_6-mer | 56,688 | 45,947 | 37041 |

*Supplementary Table 4: Correlation of $OE_{prom\_1kbps\_5\text{-}mer}$ of LPHN1, CDK9 and TRIM8 genes from chromosome 19, 9 and 10 respectively.*

| ChrA | Gene A | ChrB | Gene B | Corr 1kb_5-mer |
|---|---|---|---|---|
| 19 | LPHN1 | 9 | CDK9 | 0.905 |
| 19 | LPHN1 | 10 | TRIM8 | 0.903 |
| 10 | TRIM8 | 9 | CDK9 | 0.910 |

*Supplementary Table 5: Common transcription factors of genes having high correlation and common transcription factors. Column 1 and 2 shows the gene sets and the common transcription factors between them. The consensus from JASPAR for the respective transcription factors is shown in column 3. Column 4 records the $OE_{prom\_1kb\_5\text{-}mer}$ of genes.*

| Genes Connected | TF | Consensus Motif | Promoter Motif: OE |
|---|---|---|---|

<table>
<tr>
<td>LPHN1,CDK9,TRIM8</td>
<td>SP1</td>
<td>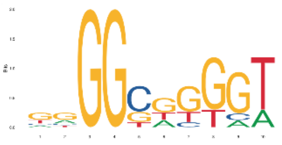</td>
<td>
<table>
<tr><th>Motif Pair</th><th>ADGRL1</th><th>CDK9</th><th>TRIM8</th></tr>
<tr><td>GGGGC</td><td>17.6</td><td>26.59</td><td>20.83</td></tr>
<tr><td>GGCGC</td><td>16.34</td><td>23.95</td><td>23.38</td></tr>
<tr><td>CGGCG</td><td>16.33</td><td>29.22</td><td>22.11</td></tr>
<tr><td>GGCGG</td><td>16.15</td><td>51.61</td><td>23.49</td></tr>
<tr><td>CCGGG</td><td>14.36</td><td>21.28</td><td>25.97</td></tr>
<tr><td>GCGGC</td><td>14.35</td><td>41.19</td><td>29.91</td></tr>
<tr><td>GGGCG</td><td>14.32</td><td>30.55</td><td>20.83</td></tr>
<tr><td>CGGGG</td><td>13.68</td><td>23.91</td><td>27.21</td></tr>
<tr><td>GCCGG</td><td>11.74</td><td>15.96</td><td>20.76</td></tr>
<tr><td>GGGGG</td><td>11.6</td><td>21.27</td><td>20.54</td></tr>
<tr><td>GCGGG</td><td>11.14</td><td>25.19</td><td>20.8</td></tr>
<tr><td>GCCCG</td><td>9.15</td><td>23.92</td><td>20.79</td></tr>
<tr><td>GGAGG</td><td>8.89</td><td>26.11</td><td>20.29</td></tr>
<tr><td>GGGCC</td><td>8.51</td><td>19.93</td><td>14.29</td></tr>
<tr><td>GGGAG</td><td>8.31</td><td>14.07</td><td>19.37</td></tr>
<tr><td>CGCGG</td><td>7.83</td><td>17.29</td><td>24.68</td></tr>
<tr><td>GGAGC</td><td>7.73</td><td>7.52</td><td>9.24</td></tr>
<tr><td>AGGCG</td><td>7.71</td><td>10.3</td><td>5.54</td></tr>
<tr><td>GGCCG</td><td>7.2</td><td>23.91</td><td>23.37</td></tr>
<tr><td>CTCCG</td><td>7.13</td><td>14.97</td><td>7.39</td></tr>
<tr><td>GCGCG</td><td>6.53</td><td>10.66</td><td>22.08</td></tr>
</table>
</td>
</tr>
<tr>
<td>LPHN1,CDK9,TRIM8</td>
<td>TFAP2A</td>
<td>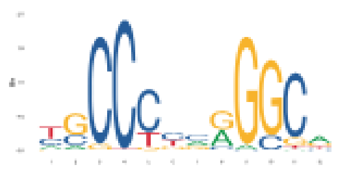</td>
<td>
<table>
<tr><th>Motif Pair</th><th>ADGRL1</th><th>CDK9</th><th>TRIM8</th></tr>
<tr><td>GGGGC</td><td>17.6</td><td>26.59</td><td>20.83</td></tr>
<tr><td>GGCGC</td><td>16.34</td><td>23.95</td><td>23.38</td></tr>
<tr><td>CGGCG</td><td>16.33</td><td>29.22</td><td>22.11</td></tr>
<tr><td>GGCGG</td><td>16.15</td><td>51.61</td><td>23.49</td></tr>
<tr><td>CCGGG</td><td>14.36</td><td>21.28</td><td>25.97</td></tr>
<tr><td>GCGGC</td><td>14.35</td><td>41.19</td><td>29.91</td></tr>
<tr><td>GGGCG</td><td>14.32</td><td>30.55</td><td>20.83</td></tr>
<tr><td>CGGGG</td><td>13.68</td><td>23.91</td><td>27.21</td></tr>
<tr><td>GCCGG</td><td>11.74</td><td>15.96</td><td>20.76</td></tr>
<tr><td>GGGGG</td><td>11.6</td><td>21.27</td><td>20.54</td></tr>
<tr><td>GCGGG</td><td>11.14</td><td>25.19</td><td>20.8</td></tr>
<tr><td>GCCCG</td><td>9.15</td><td>23.92</td><td>20.79</td></tr>
<tr><td>GGAGG</td><td>8.89</td><td>26.11</td><td>20.29</td></tr>
<tr><td>GGGCC</td><td>8.51</td><td>19.93</td><td>14.29</td></tr>
<tr><td>GGGAG</td><td>8.31</td><td>14.07</td><td>19.37</td></tr>
<tr><td>CGCGG</td><td>7.83</td><td>17.29</td><td>24.68</td></tr>
<tr><td>GGAGC</td><td>7.73</td><td>7.52</td><td>9.24</td></tr>
</table>
</td>
</tr>
</table>

| | | | |
|---|---|---|---|
| | | | AGGCG 7.71 10.3 5.54<br>**GGCCG** 7.2 23.91 23.37<br>CTCCG 7.13 14.97 7.39<br>GCGCG 6.53 10.66 22.08 |
| LPHN1,TRIM8 | **E2F1** | 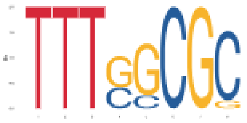 | Motif<br>Pair LPHN1 TRIM8 CDK9<br>TTTTC 3.20 1.32 0<br>TTTCC 2.97 0.93 0.47<br>TGCGC 2.98 6.46 0.64 |
| TRIM8,CDK9 | **HINFP** | 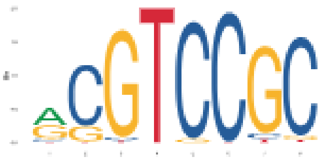 | Motif<br>Pair CDK9 TRIM8 LPHN1<br>ACCCC 4.7 3.69 0.58<br>TCCTC 6.57 5.23 1.63<br>TCCGG 6.56 6.46 2.98<br>AGGGG 8.44 8.32 4.76<br>GTCCG 1.87 1.84 0.6 |
| LPHN1,CDK9 | **RELA** | 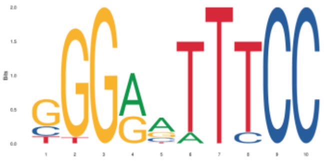 | Motif<br>Pair LPHN1 CDK9 TRIM8<br>TTCCC 2.16 1.32 0.00 |

*Supplementary Table 6: Comparison of number of genes annotated according to different reference assemblies.*

| Chromosome | T2T | GRCh38 | GRCh37 |
|---|---|---|---|
| Chr18 | 1022 | 983 | 717 |
| Chr19 | 2531 | 2492 | 2248 |

*Supplementary Table 7: Comparison of gene-pair correlation inter chromosome 18 and chromosome 19 in Hi-C vs pcHi-C vs all*

| Resolution | 1kb_5-mer | 1kb_6-mer |
|---|---|---|
| Gene pairs with motif vector correlation >= 0.5 | 9.92% | 0.47% |
| Gene pairs in Hi-C with motif vector correlation >= 0.5 | 14.38% | 1.01% |
| Gene pairs in pcHi-C with motif vector correlation >= 0.5 | 22.80% | 0.85% |

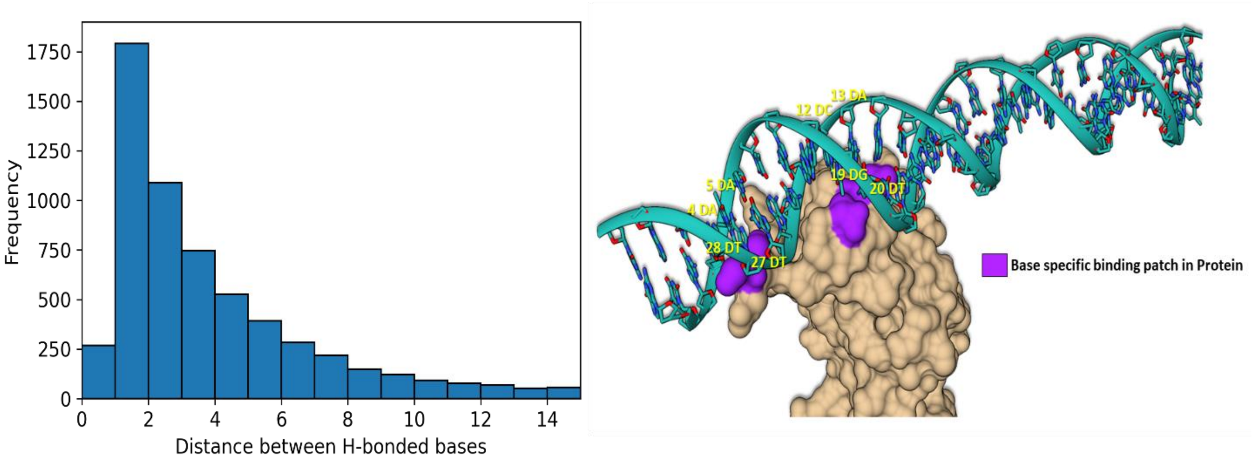


a. b.

*Supplementary Figure 1a: Frequency distribution of sequence distances between hydrogen bonded nucleotides in an NR set of 285 DNA bound protein complexes [30]. Figure 1b. An example where the distance between two DNA bases involved in a specific interaction is more than 6bps (PDB ID: 4jl3). The protein is shown in surface representation and the DNA is in ribbons and sticks. The specific bases that make hydrogen bonds with the proteins are labelled (in yellow). Notice 1) Only ~15% of the hydrogen bonds are separated by >6 base pairs; 2) Hydrogen bonds that are separated by >6 base pairs interact with amino acids of two distinct patches on the surface of the protein (shown in purple).*

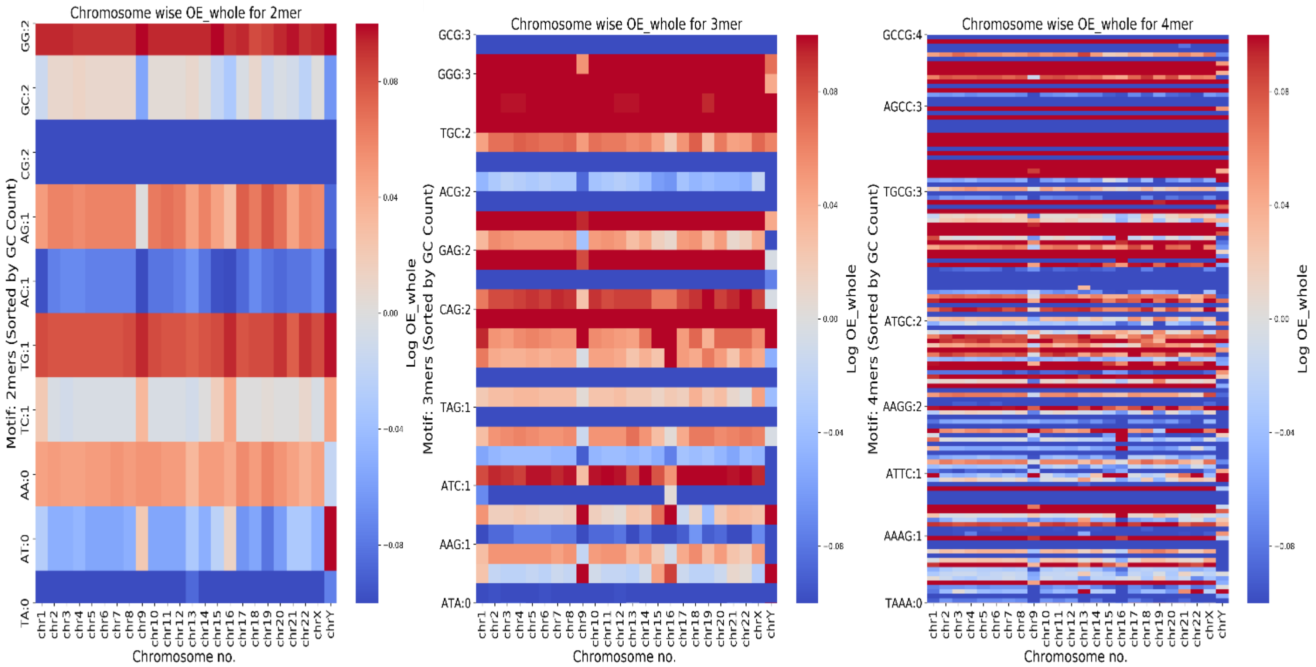


a. b. c.

*Supplementary Figure 2: Heat maps of $OE_{whole}$ ratios for motif size 2-4 (panels a, b and c respectively) across the different chromosomes shown in log scale. The log(OE) ratios are coloured red through to blue (colour legend) where red and blue indicate values of > 1 and < 1 respectively. The darker the shade of the red (or blue) the higher (or lower) the ratio. The motifs are arranged according to GC content, which increases going from bottom to top. In panels b and c, only a few representative motifs (and their reverse complements) are labelled. Shown alongside the motif labels are their GC contents.*

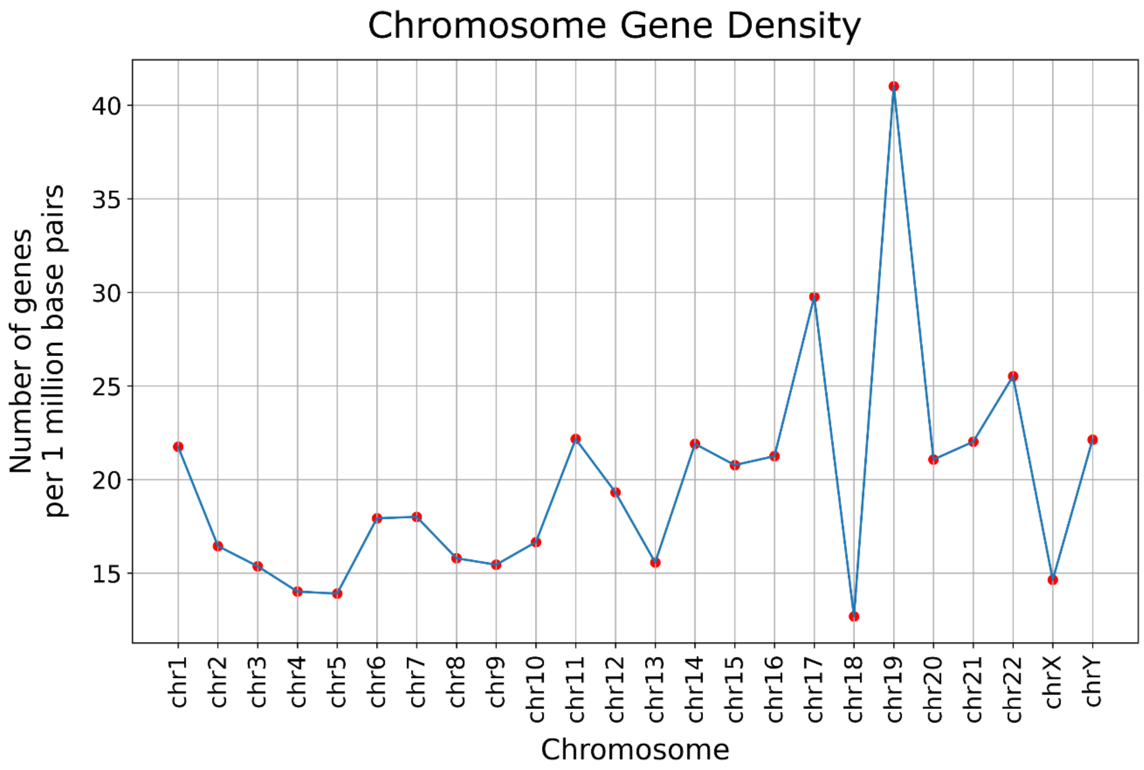


*Supplementary Figure 3: Chromosome-wise gene density plot. The number of genes per million bases is shown for the different chromosomes.*

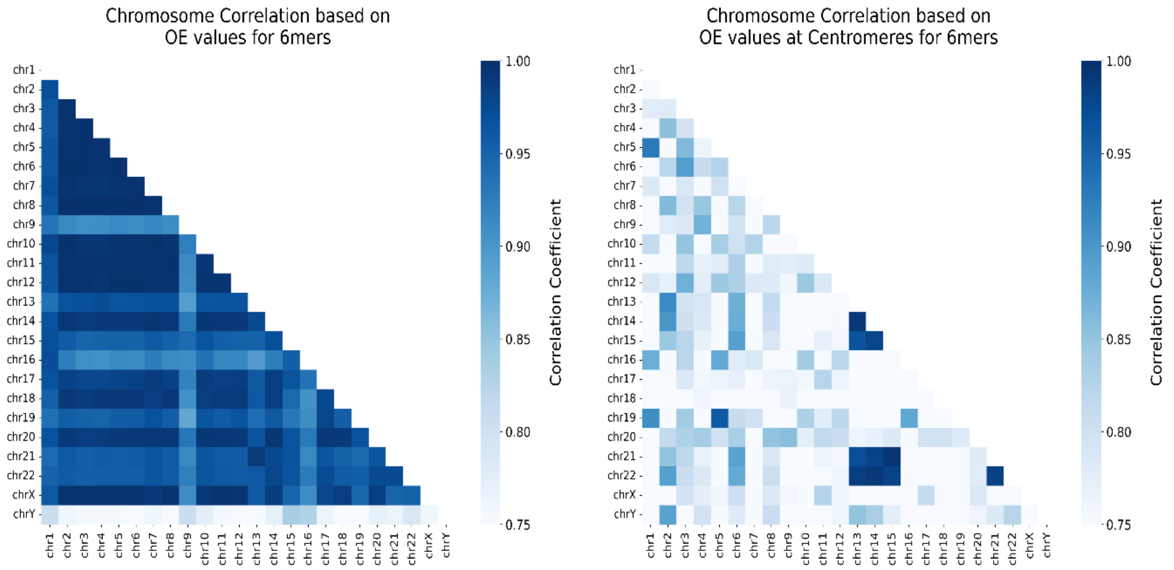


a. b.

*Supplementary Figure 4: Heat maps of the Pearson's correlation coefficient based on 6mer motif vectors in the whole chromosome (4a) and centromeric regions (4b). The x- and y- axes represent the chromosome pairs. The correlation coefficients are coloured white through to blue. The darker the shade of the blue the higher the correlation.*

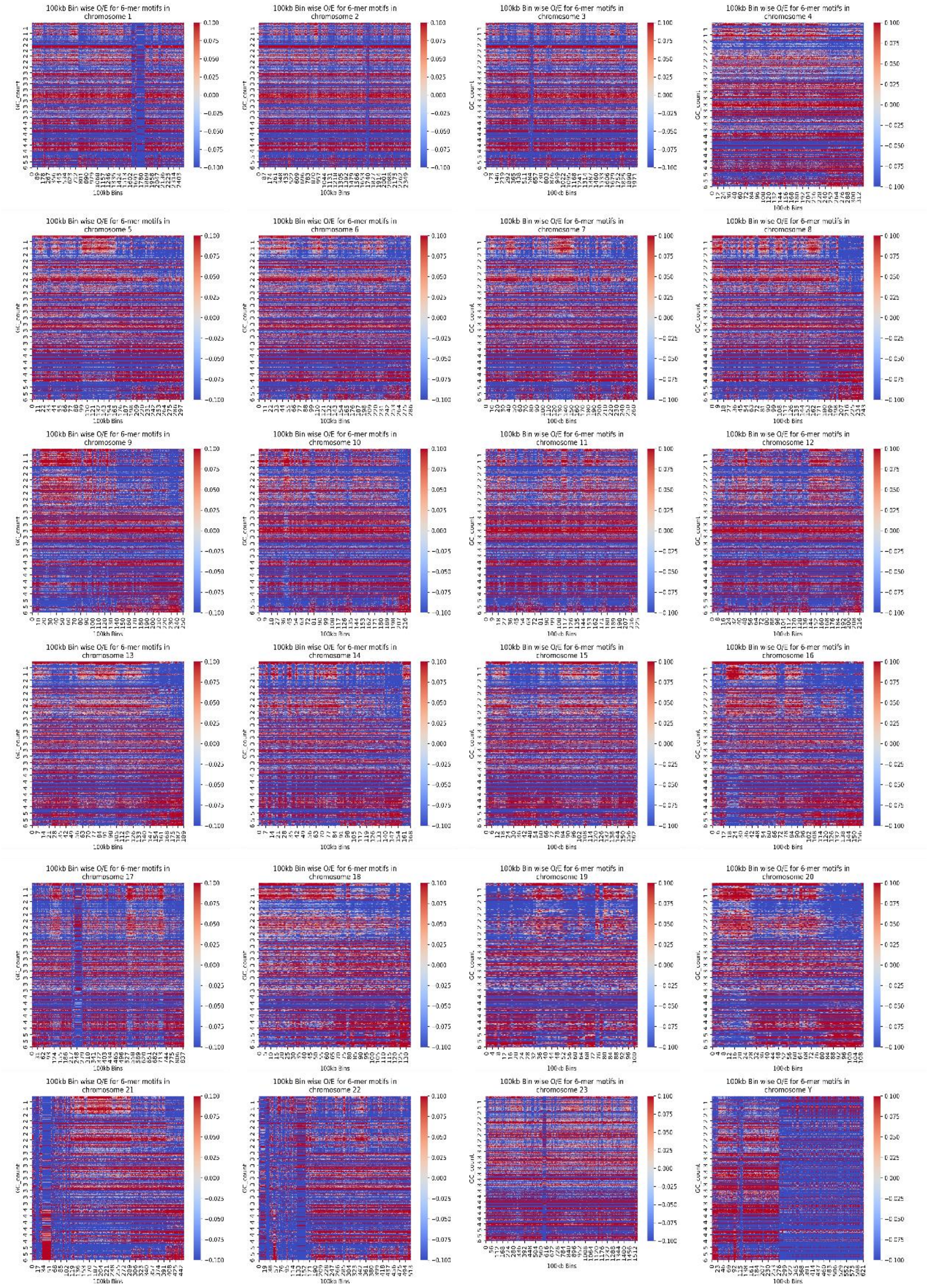


*Supplementary Figure 5: Heat maps of the log $OE_{100kb}$ for all actual chromosome sequences, for motif size 6 mers. The log(OE) ratios are coloured red through to blue (colour legend) where red and blue indicate values of > 1 and < 1 respectively. The darker the shade of the red (or blue) the higher (or lower) the ratio. The 512 and 2080 motifs for 5-mers and 6-mers respectively are arranged according to GC content, which increases going from bottom to top. On y-axis the GC-content of the motifs are labelled.*

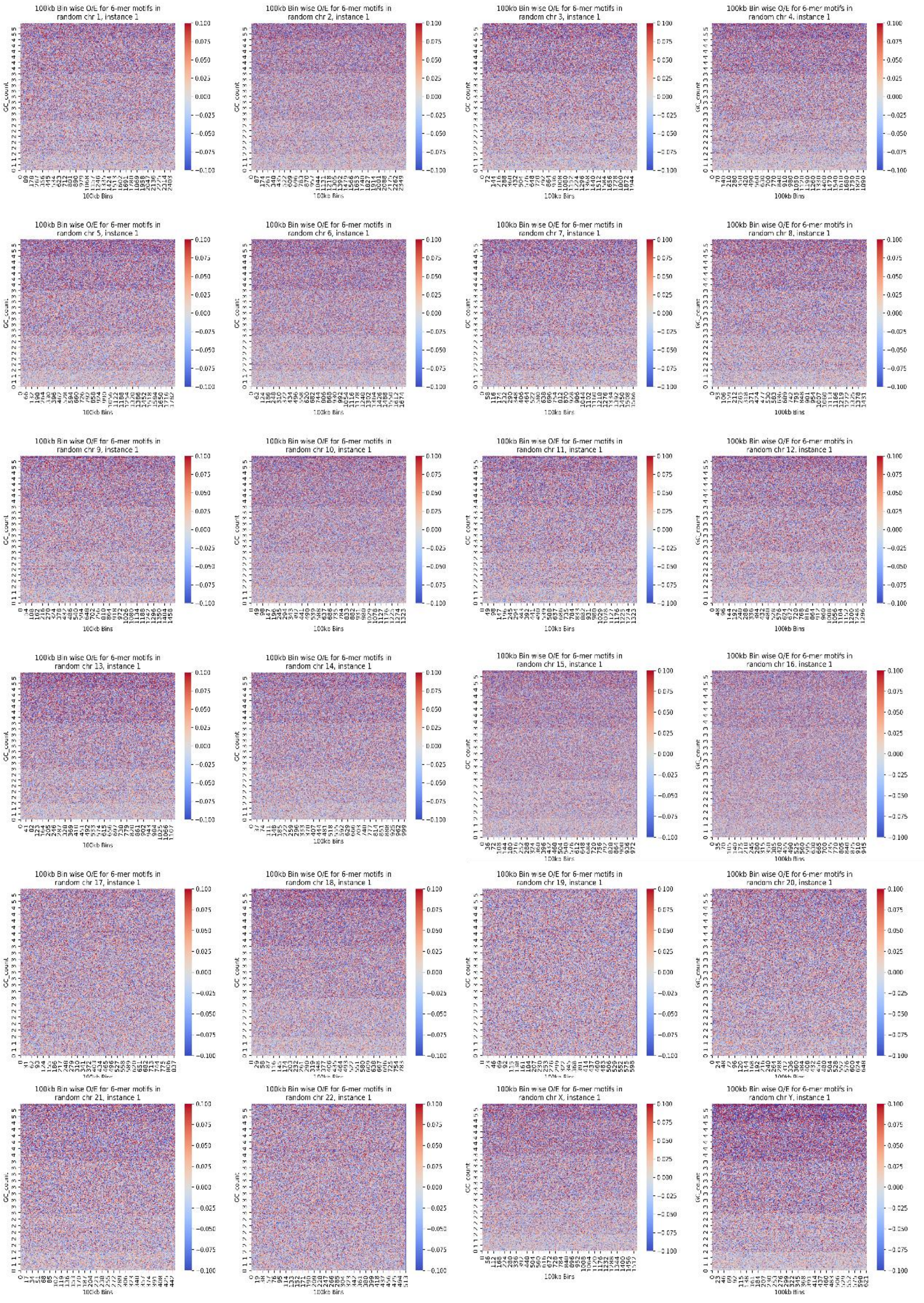


*Supplementary Figure* *<u>6</u>*: *Heat maps of the log* $OE_{100kb}$ *for all random chromosome sequences, for motif size 6 mers. The log(OE) ratios are coloured red through to blue (colour legend) where red and blue indicate values of > 1 and < 1 respectively. The darker the shade of the red (or blue) the higher (or lower) the ratio. The 512 and 2080 motifs for 5-mers and 6-mers respectively are arranged according to GC content, which increases going from bottom to top. On y-axis the GC-content of the motifs are labelled.*

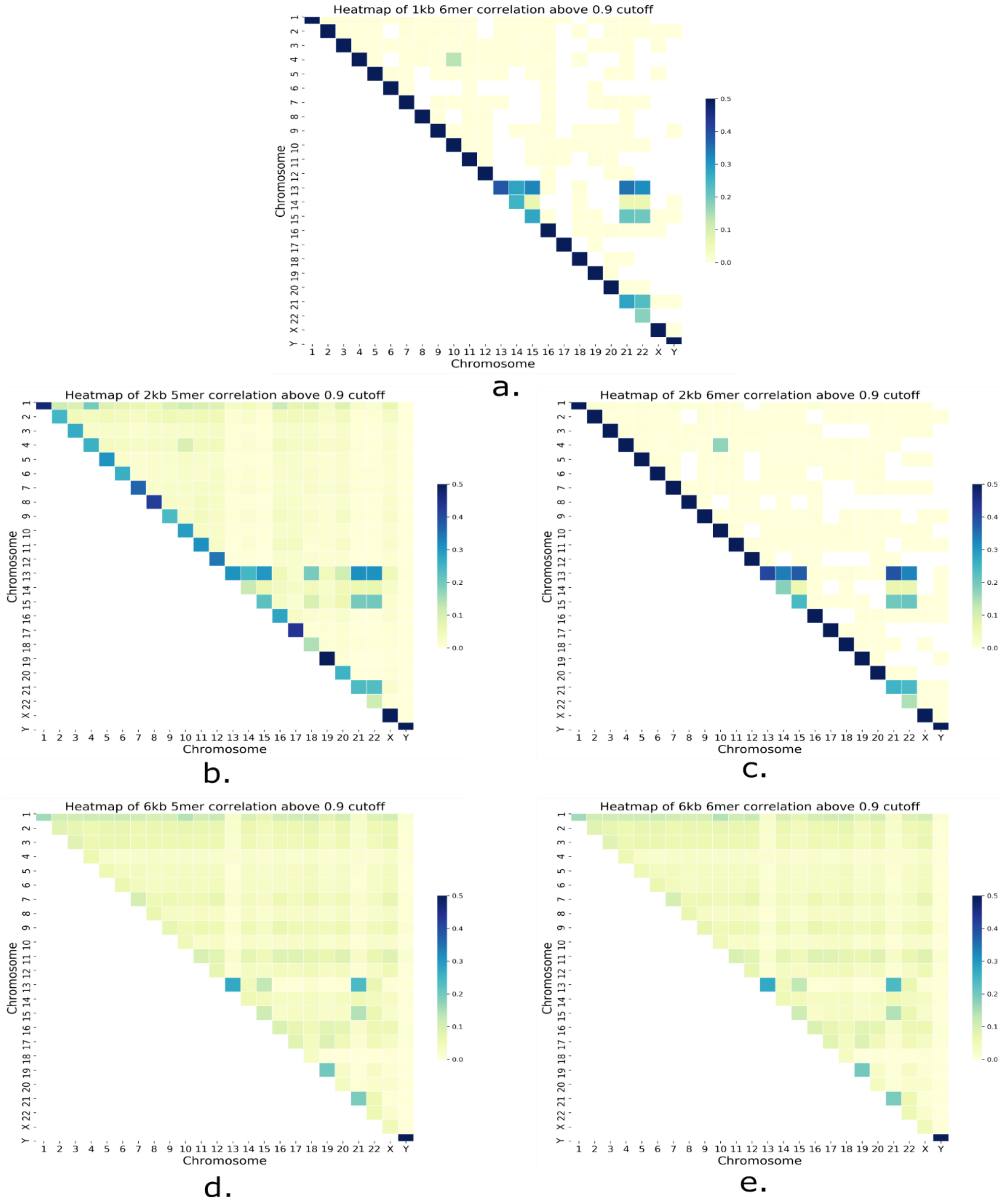


*Supplementary Figure 7: An inter-chromosome motif vector correlation map of frequency of gene pairs with correlation coefficients > 0.9 for 5 mers with promoter sizes of 1,2 and 6kbps for motif sizes 5 and 6 (a-e). The correlations are colour coded on a white-yellow-green-blue gradient. The white end represents no correlation while the darker the shade of blue the larger are the number of correlated gene pairs.*

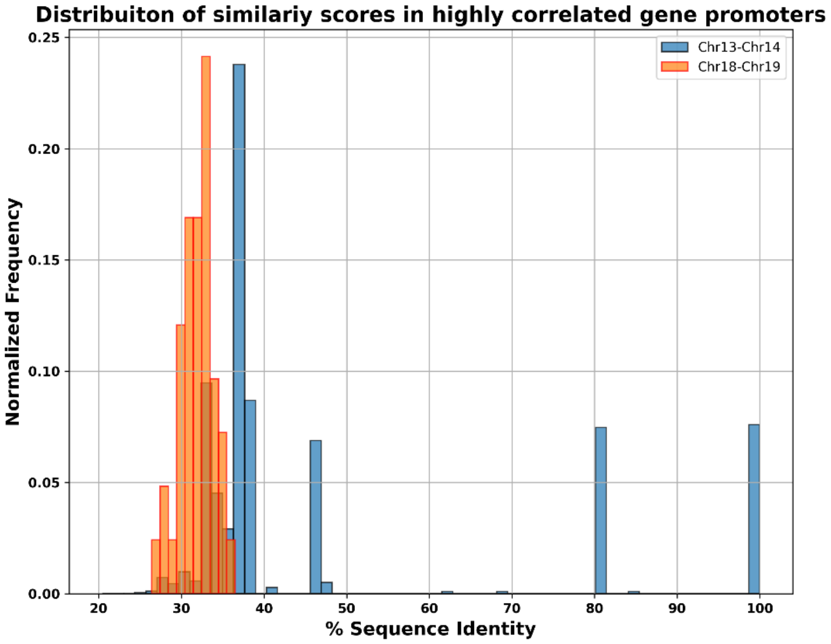


*Supplementary Figure 8: Distribution of sequence similarity in gene promoters having high correlation in chromosome pair 13-14 (blue) and 18-19 (orange). The x-axis represents the percentage sequence identity in the promoters and the y-axis represents the normalized frequency.*

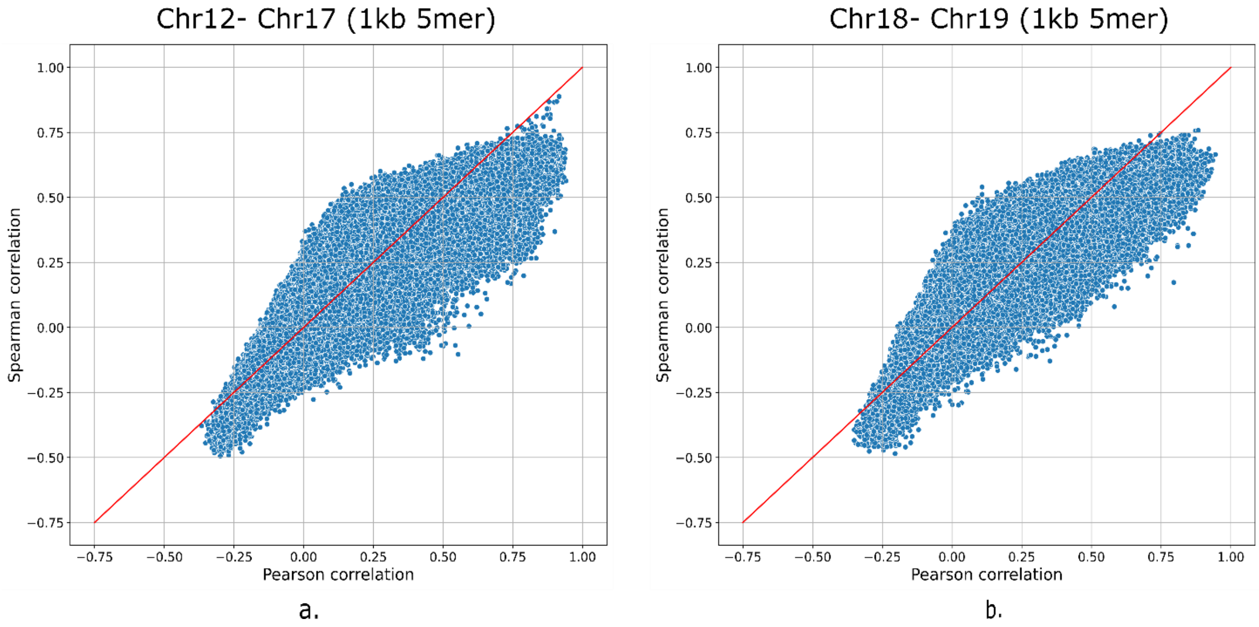


*Supplementary Figure 9: Comparison of Spearman vs Pearson's correlation coefficient for motif vector correlations of all gene promoters. The x-axis represents Pearson's correlation score and the y-axis represents Spearman's score. Figure a and b show the comparisons for chromosome pairs 12-17 and 18-19 respectively.*

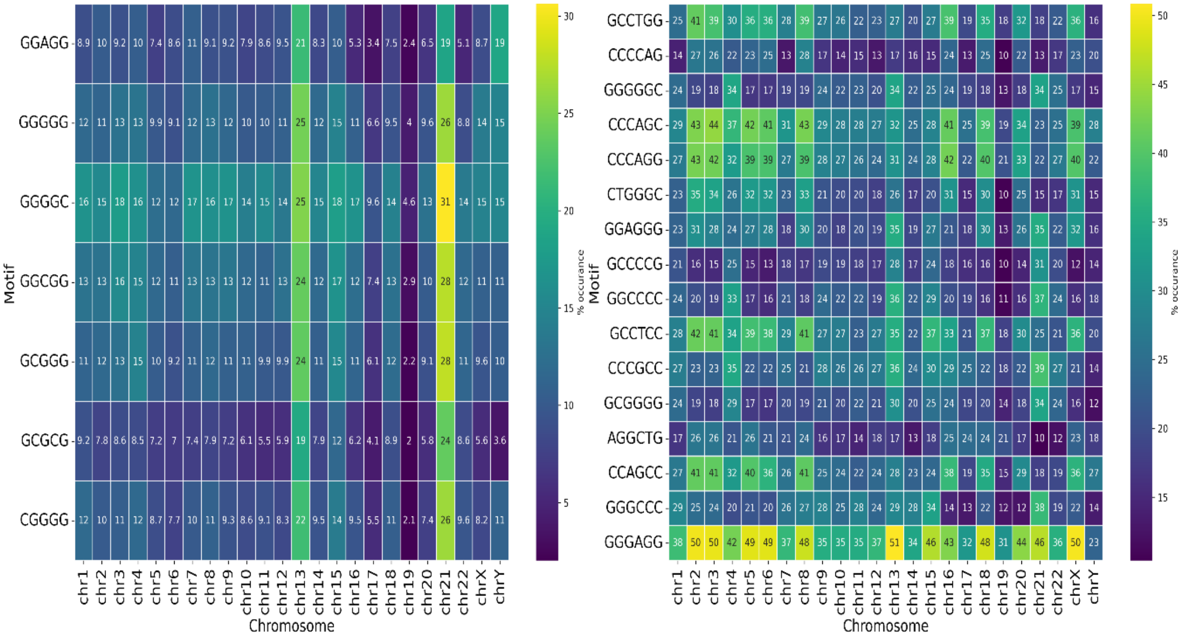


a. b.

*Supplementary Figure 10: a) Seven and b) sixteen 6 5- and 6-mers respectively that have an abundance of OE>10 in at least 1% of promoter proximal control regions in every single chromosome. The heatmap represents the percentage of genes in the different chromosomes where the motif has OE >10. The lighter the shade the larger the number of promotor proximal control regions the motif is present in. The numerical values in the cells represent the percentage of the total genes the motifs are present in that chromosome.*

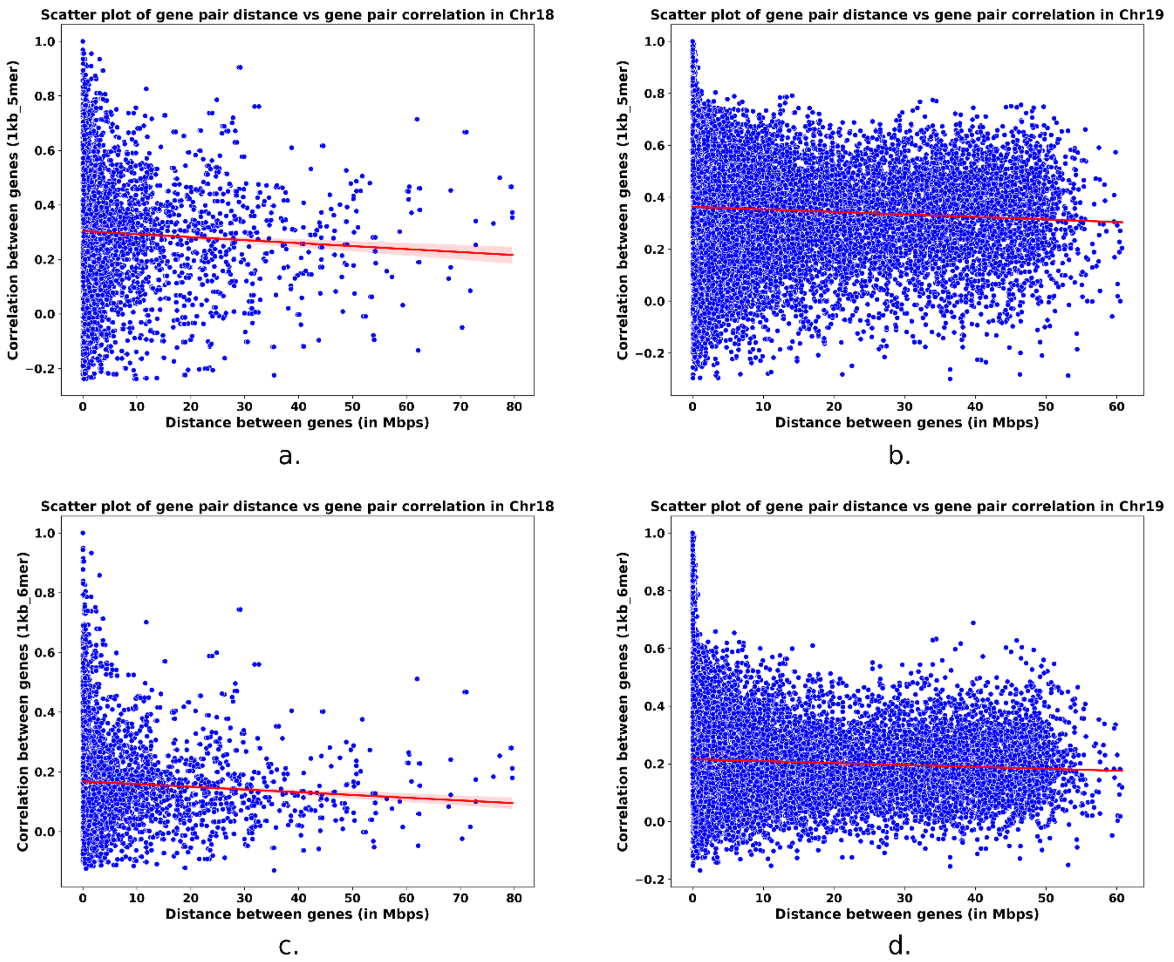


*Supplementary Figure 11: Comparison of distance between genes and motif vector correlations. The y-axis represents the correlation coefficient and the x-axis represents the distance between gene pairs in (Mbps). Panels a and b show motif vector correlations at 1kb 5-mer for gene pair in chromosomes 18 and 19 respectively. Panels c and d show the correlations at 1kb 6mer resolutions in the same order.*

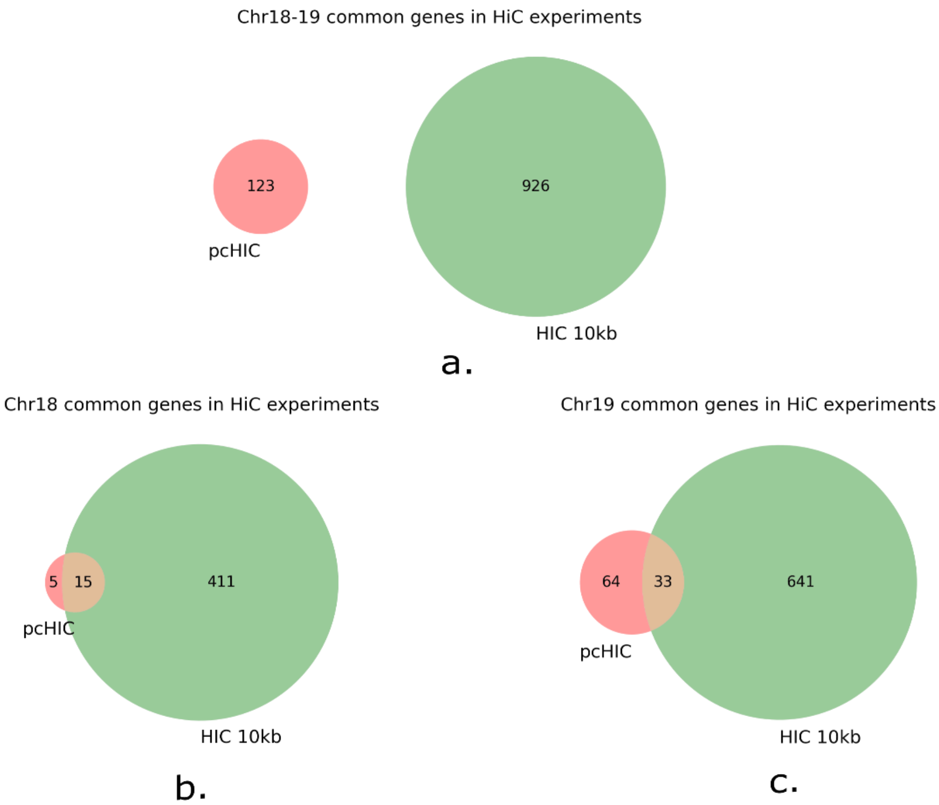


*Supplementary Figure 12: Overlap of gene/gene-pairs in Hi-C vs pcHi-C data. Overlap between all the gene-pairs [a]. The overlap between the unique genes involved in different Hi-C contacts in Chr18[b] and Chr19[c]. Green colour represents Hi-C capture at 10kbps resolution and peach colour represents pcHi-C contacts.*

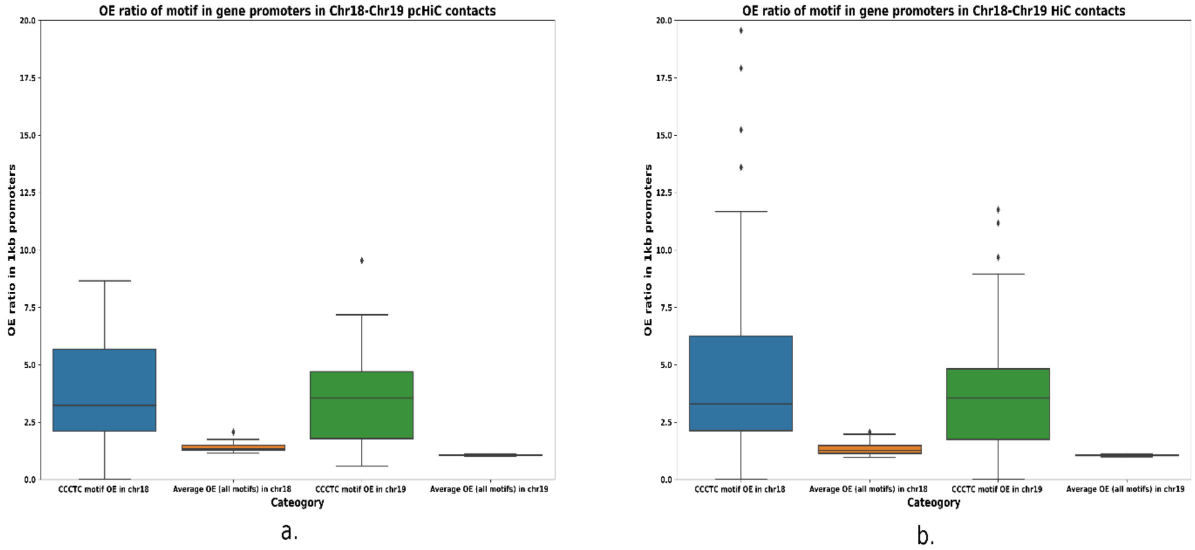


*Supplementary Figure 13: Distribution of $OE_{1kbps\_5\text{-}mer}$ for the CTCF binding motifs vs other motifs in gene promoters with Hi-C and pcHi-C contacts between chromosome 18-19 Hi-C. Panels a and b represent the distribution of genes from pcHi-C and Hi-C experiments respectively. Blue and green colour box plot represents the distribution of the CCCTC motif in chromosome 18 and chromosome 19 respectively. Orange and black coloured boxes represent the distribution of other motifs in chromosome 18 and chromosome 19 respectively.*
